# scRNA-seq reveals the diversity of the developing cardiac cell lineage and molecular building blocks of the primary pacemaker

**DOI:** 10.1101/2023.06.26.546508

**Authors:** Karim Abu Nahia, Agata Sulej, Maciej Migdał, Natalia Ochocka, Richard Ho, Bożena Kamińska, Marcin Zagorski, Cecilia L. Winata

## Abstract

The heart is comprised of a variety of specialized cell types that work in unison to maintain blood flow. Here we utilized scRNA-seq analysis to delineate the diversity of cardiac cell types in the zebrafish. With the growing use of the zebrafish to model human heart biology, a deeper insight into its complex cellular composition is critical for a better understanding of heart function, development, and associated malformations. We present a high resolution atlas of zebrafish heart single cells transcriptomics, consisting of over 50 000 cells representing the building blocks of the zebrafish heart at 48 and 72 hpf. We defined 18 discrete cell populations comprising major cell lineages and sublineages of the developing heart. We pinpointed a population of cells likely to be the primary pacemaker and identified the transcriptome profile defining this critical cell type. Our analyses identified two genes, *atp1b3b* and *colec10*, which were enriched in the sinoatrial pacemaker cells. CRISPR/Cas9-mediated knockout of these two genes significantly reduced heart rate which is accompanied by arrhythmia or morphological defects, suggesting their novel function in cardiac development and conduction. Additionally, we describe other subpopulations of cardiac cell lineages, including the endothelial and neural cells, whose expression profiles we provide as a resource for further investigations into the cellular and molecular mechanisms of this organ.

## Introduction

Essential components of the heart ensure its life-sustaining activity [1]. Specialized cell types constitute the contractile myocardium of the atria and ventricles, while cells of the cardiac conduction system and the autonomous nervous system innervates the heart tissues and coordinate rhythmic contractions of the heart chambers [2]. The two pacemakers, sinoatrial (SA) and atrioventricular (AV) nodes, spontaneously generate electrical impulses driving heart contraction [3]. Endothelial cells form the inner endocardial lining of the heart lumen [4], a subset of which are further specialized to form the heart valves [5], while a separate population makes up the coronary blood vessels that supply oxygen for myocardial contraction [6]. The epicardium provides a protective layer surrounding the heart muscles [7]. Besides these main sublineages, other cell types, mainly fibroblasts and mesenchyme, provide the structural matrix of the organ and play multiple roles in physiological processes related to cardiac function and regeneration [8]. The heart also performs other functions such as endocrine and iron homeostasis (reviewed in [9,10]) which are established though less well described.

The core genetic program and stepwise morphogenesis involved in the development of the heart is largely conserved across metazoans [11]. Cells making up the heart arise from a pool of common mesodermal progenitors which are specified to the various major lineages. In the zebrafish, these progenitors can be detected by 12 hours post-fertilization (hpf) by the expression of *myl7* and *nkx2.5*. Subpopulations of myocardial progenitors could be further distinguished between atrial (expressing *myh6*) and ventricular (expressing *myh7*) myocardium [12]. Concurrently, endocardial progenitors denoted by the expression of *kdrl* and *cdh5* could be found anterior to the myocardial progenitors [13]. As the myocardial progenitors migrate to the midline to form a heart tube by 19 hpf, endocardial progenitors proceed to migrate towards the median and line the lumen of the heart tube [14]. By 22 hpf, a linear, beating heart tube is formed, which is continuously elongated by the addition of cells originating from the second heart field (SHF), a pool of late-differentiating mesodermal progenitors, extending the atrium and ventricle and forming inflow and outflow tracts [15,16]. Between 24 and 30 hpf, neural crest cells (NCCs) migrating through the pharyngeal arches contribute to the myocardium of the primitive heart tube, while a second population arriving at 80 hpf contribute to the outflow tract structures [17]. The NCCs also contribute to the cardiac peripheral nervous system [18]. Finally, the proepicardial organ forms at 48 hpf close to the atrioventricular junction and can be detected by the expression of *wt1* and *tcf21* markers. This structure gives rise to the epicardial cells which subsequently migrate and envelop the heart [7].

Although key insights into heart development and function have been derived from the zebrafish model organism [11,13], critical differences exist between the zebrafish and mammalian heart. Unlike mammals, teleosts including the zebrafish possess a heart comprising two chambers. An additional structure unique to teleosts is the bulbus arteriosus (BA), a chamber-like structure located at the outflow portion of the heart which is composed of smooth muscle and serves to absorb pressure [19]. In terms of electrophysiology, the zebrafish heart is fundamentally similar to that of humans, which enables faithful modeling of diseases of the cardiac conduction system [20,21]. Yet, differences in terms of ion channels and the types of currents involved in driving cardiac contractions exist [22]. With the growing use of the zebrafish to model human heart biology, a comprehensive knowledge of both conserved and nonconserved features between the hearts of the two organisms becomes necessary in order to more accurately translate results from the zebrafish to human.

Single cell analyses of mammalian heart have revealed surprising new insights in the discovery of new cell types and their contributions to various forms of heart disease [23–26]. The zebrafish offers a glimpse into earlier events of cardiogenesis which could provide valuable insights into the mechanism underlying the development of various cardiac cell types and specialized structures. Several single cell level analyses have been performed in zebrafish which included the heart [27–30]. However, these were either focused on selected cell types or have not reached sufficient depth to comprehensively capture and annotate cardiac cell subtypes, particularly rare cell types such as the cardiac pacemaker cells. These cells are embedded within the myocardium and play a central role in generating and propagating electrical impulses for a rhythmic heart contraction. Although previous bulk-level analyses have shed some light into the molecular mechanism of their function [31–33], their scarcity and the lack of defining morphological and molecular features continue to pose a challenge to isolate pure populations of this cell type and study them.

Here, we present a high resolution atlas of the developing zebrafish whole heart single cells transcriptomics, aiming at sufficient depth to allow discovery and annotation of cardiac cell subtypes. We delineated major cell lineages and sublineages of the zebrafish heart and distinguished a set of gene expression profiles associated with each of these populations. We uncovered new cell subpopulations within major clusters of cardiomyocytes, endocardium, neural-crest-derived, and fibroblast cells. Clustering analyses revealed two novel genes specifically enriched in the SA pacemaker, *colec10* and *atp1b3b*, which encode the collectin subfamily member 10 [34] and a subunit of the Na+/K+ ATPase beta chain proteins [35], respectively. Loss of function analyses further revealed their role in heart development and rhythm maintenance. Our study established the heterogeneity of zebrafish cardiac cell types which could serve as a valuable resource for future in-depth analyses of cell populations with higher specificity.

## Methods

### Zebrafish

Zebrafish transgenic lines *sqet31Et* [33,36,37]*, sqet33mi59BEt* [32,36,37], *Tg(myl7:mRFP)* [38], *Tg(myl7:EGFP)* [39] or *Tg(myl7:EGFP-Hsa.HRAS)^s883^* [40], were maintained in the zebrafish facility of the International Institute of Molecular and Cell Biology in Warsaw (license no. PL14656251) in line with standard procedures and ethical guidelines. Embryos were raised in egg water at 28°C, screened for fluorescence signals in the heart and staged at 48 hpf and 72 hpf based on established morphological criteria [41]. All zebrafish embryos utilized for microscopy were supplemented with 0.003% PTU (Sigma-Aldrich, P7629) in E3 medium (5 mM NaCl, 0.17 mM KCL, 0.33 mM CaCl, 0.33 mM MgSO4, pH 7.4) shortly before 24 hpf to inhibit pigmentation and ease heart rate observations [42].

### Heart extraction and single cell dissociation

Whole hearts were extracted from double transgenic individuals [*sqet31Et* x *Tg(myl7:mRFP)* and *sqet33mi59BEt* x *Tg(myl7:mRFP)*]. The term “pseudo-replicates” was used to denote samples of two different genotypic backgrounds (*sqet31Et* and *sqet33mi59BEt*) at the same developmental stage. Embryos were anesthetized with Tricaine (0.16 mg/ml in egg water) and large-scale extraction was performed according to a previously published protocol, with minor adjustments [43]. Hearts were manually separated from remaining tissue under a fluorescence stereomicroscope and collected into 0,5 ml of EDM (L-15/10% FBS). Single cell heart suspension was obtained according to Bresciani et al. [44]. Ultimately, cells were resuspended by gentle pipetting, loaded on the cell counting chamber, visually inspected under the microscope (Zeiss Apotome), and quantified by Countess automated cell counter (Invitrogen) (Fig. 1A, Supplementary Fig. 1A).

**Figure 1.**
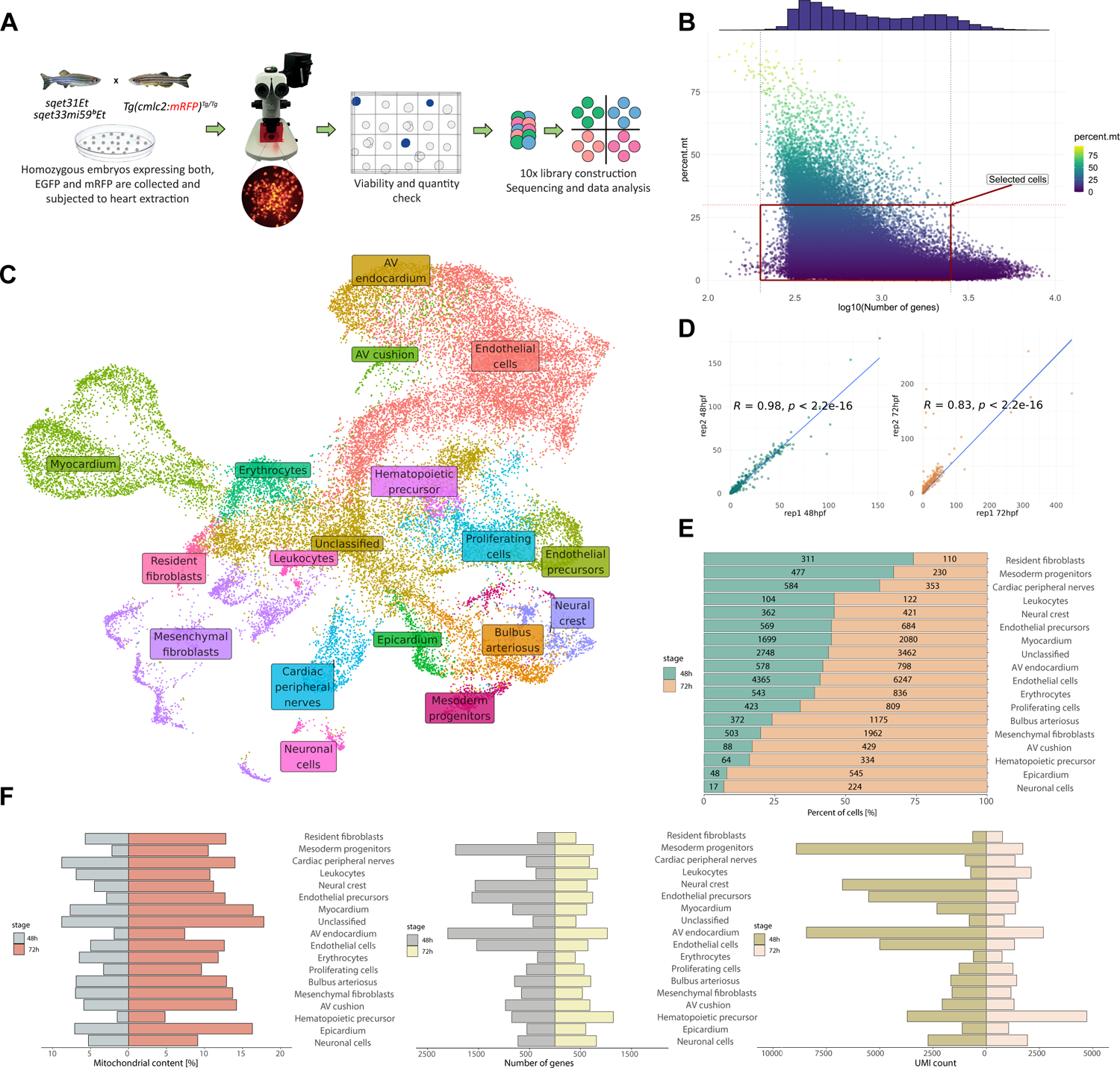
Single-cell RNA sequencing revealed 18 distinct cell subtypes of the embryonic zebrafish heart. **A** Schematic overview of experimental design and workflow. **B** Quality parameters of all cells derived from 48 and 72 hpf pseudo-replicates showing the number of expressed genes and mitochondrial gene content. Cells within the red frame were included in the downstream analysis. **C** Integrated UMAP projection depicting cell clusters that constitute the embryonic zebrafish heart. **D** Pearson correlation between developmental pseudo-replicates at both 48 hpf and 72 hpf. **E** Visual representation of cluster composition expressed as a percentage indicating the number of cells contributing to each cluster depending on developmental stage. **F** General quality metrics reflecting the percent of mitochondrial genes, the total number of genes, and UMI expressed in each cell cluster at a particular developmental stage.

### Library prep and sequencing

After viability and quantity check, dissociated cells derived from embryonic zebrafish hearts at 48 hpf and 72 hpf were loaded on 10x Chromium Controller Chip G (10x Genomics) and processed according to the Chromium Next GEM Single Cell 3’ Reagent Kits v.3.1 targeting 10 000 cells. Resulting libraries were verified by High-Sensitivity DNA Kit (Agilent Technologies) on a 2100 Bioanalyzer (Agilent Technologies) and Qubit Fluorometer using Qubit dsDNA High-Sensitivity Assay Kit (Invitrogen). Final libraries were quantified with KAPA Library Quantification Kit Illumina® Platforms (Kapa Biosystems/Roche), followed by paired-end sequencing (read 1 - 28 cycles, i7 index - 8 cycles, i5 index - 0 cycles, read 2 - 91 cycles) performed with Nextseq 500 (Illumina).

### QC, filtering and clustering

Obtained raw sequencing data (BCL files) were demultiplexed and transformed into fastq files using 10x Cellranger pipeline version 3.1.0 [45] and bcl2fastq v2.19.0.316 (Illumina). The resulting matrices were loaded into the Seurat package v4.0.1 [46] for R v4.0.2 [47] applying standard quality control, normalization and analysis steps unless otherwise specified in the description. Briefly, sequencing reads were mapped to the zebrafish reference genome GRCz11 (Ensembl release 100) extended with additional EGFP and mRFP sequences. Due to overlapping annotations, reads mapped to non-polyA transcripts and protein-coding genes may be flagged as multi-mapped, and in consequence not count in the 10x pipeline, respective annotation GTF file was pre-filtered to retain only protein-coding genes (*cellranger mkgtf -- attribute=gene_biotype:protein_coding*). The estimated number of cells identified by the Cellranger pipeline was 52,447 (Supplementary Table 1). In order to dispose of potential empty droplets, low quality cells and possible multiplets, droplets with a very small (< 200) and very high (> 2500) library size were removed. Standard scRNA-seq filtering workflows exclude cells with a high ratio of reads from mitochondrial genome transcripts, which may indicate potential cell membrane rupture and dissociation-based damage. This filter is often set at 5-10% [48], however, the heart as muscle tissue is composed of diverse cell populations including cardiomyocytes which require high energy demand leading to potentially very high mitochondrial gene content per cell. Therefore, to retain cardiomyocytes while removing low-quality cells, we chose a threshold of 30% mitochondrial gene content. In addition to above mentioned thresholds, we also decided to remove cells containing more than 10% of hemoglobin reads. All together, to keep high quality cells the following thresholds were applied: 200 < number of genes < 2500 & mitochondrial gene content < 30% & hemoglobin gene content < 10%. An increasing number of targeted cells is inherently associated with higher multiplet rate. To maximize the removal of possible doublets from the dataset, in addition to filtering out cells with very high gene content (> 2500) we applied the DoubletFinder [49] approach to exclude extra heterotypic doublets derived from transcriptionally distinct cells (Supplementary Table 1). After applying all filtering steps, a total of 34 676 cells were retained for further analysis (Supplementary Table 1). Following the Seurat workflow, the resulting gene expression matrices were normalized according to “SCT transform” using “glmGamPoi” method and the default number of cells and variable genes [50]. The scaled matrices were dimensionally reduced by PCA and Uniform Manifold Approximation and Projection (UMAP) embedding followed by neighbor finding and clustering according to default Seurat workflow. In each, 30 principal components were used.

### Integration of single-cell datasets

Having two biological pseudo-replicates for each timepoint (48 hpf and 72 hpf, respectively), data were integrated using the Seurat integration approach applying default parameters [51]. 3 000 integrated anchors were identified and used as an input to create an integrated dataset as implemented in Seurat. Cells were clustered based on KNN graph based approach and Louvian algorithm for modularity optimization with the resolution parameter ranging from 0.5 to 12. Ultimately resolution was set to 1.5. Clustering results were visualized by UMAP. Finally, we used the built-in default Seurat *FindAllMarkers* and *FindMarkers* functions to identify differentially expressed genes in each cluster. As a background, the entire dataset was used except for differential expression analysis between myocardial subclusters that have been compared internally. Functional Gene Ontology (GO) analysis was performed online (http://geneontology.org/) using PANTHER classification system [52] according to the default parameters (Fischer’s exact test and FDR as correction) or ClusterProfiler v. 3.17.3 [53]. Top 100 differentially upregulated genes were used as an input. However, many gene products involved in pacemaker development in zebrafish are not comprehensively captured by GO [54], therefore, the Ensembl IDs of all upregulated genes (adj. p-value <0.05) from Sinoatrial CMs cluster were converted to human orthologs and searched against *Homo sapiens.* All subsequent enrichment plots were generated in ggplot2 package using zebrafish ensembl IDs. The single cell transcriptome data can be accessed http://zfcardioscape.iimcb.gov.pl. Interactome map between the cardiac cell clusters was constructed using the DanioTalk tool [55]. The tool provides zebrafish-focused annotations based on physical interaction data of the zebrafish proteome and hence provided significantly higher number of ligand-receptor interactions compared to others built on mammalian data. All sequencing data have been deposited in the GEO database under accession number GSE234216.

### Spatial correlation analysis

The TOMO-seq dataset [31] was downloaded and used as spatial reference. Genes that had a maximum read count of less than 20 were first removed. We then smooth the data with local regression method (LOESS, span α=0.15) and calculate fold changes for all genes against the average for that gene across all sections. The Pearson correlation coefficient between each scRNA-seq clusters and each section is calculated. The spatial correlations (ρ) between the scRNA-seq clusters and the sections [31] were calculated by applying the following steps: (1) Ensembl gene IDs were used to match the data from the clusters and the sections, (2) in the cluster data, the log2FC were converted to fold change values, (3) if resulting value for a given gene ID was zero either for clusters or sections, these gene ID was ignored, (4) the Pearson correlation coefficient ρ(*x*) between the cluster and each section *x* was estimated. The average number of compared genes for all sections was 254. As a proxy for assessing quality of comparison, we investigated the number of genes that were compared for estimation of ρ for each section with respect to this average value. The sections from 2nd to 36th had number of compared genes between 0.8 and 1.2 of the average value, and only sections 37th to 39th had 0.2 of the average value. Section IDs are as in [31].

### Zebrafish F0 knockout of candidate genes using CRISPR/Cas9

The sgRNA sequences were designed utilizing chopchop [56] with default settings for CRISPR/Cas9 knock-out method. Based on previously established protocol [57], sgRNAs were designed to target three different loci per gene. The following exons were targeted: 2nd, 4th and 5th as well as 2nd, 3rd and 6th for *atp1b3b* and *colec10*, respectively (Figure 4B, Supplementary Table 7). All selected sgRNAs were specific to the targeted gene showing no mismatches and off-targets according to chopchop (Supplementary Table 7). Synthetic sgRNAs were ordered from Synthego at 1.5 nmol scales and dissolved in 15 μl of 1X TE buffer (10 mM Tris-Hcl, 1 mM EDTA, pH = 8) to reach 100 μM stock concentration. Sequences and all quality metrics are provided in Supplementary Table 7. Cas9 protein was ordered from IDT (Alt-R™ S.p. Cas9 Nuclease V3, catalog number 1081059). Based on [58], a pre-assembled complex made up of Cas9 and gRNA results in greater efficiency than the coinjection of Cas9 and gRNA. Therefore, one day prior to scheduled injections, individual sgRNAs were mixed in equal volumes and adjusted to 19 μM sgRNA mix concentration. sgRNA mixes were combined with previously diluted Cas9 protein in Cas9 buffer (20 mM Tris-HCl, 600 mM KCl, 20% glycerol [59]) to reach a final concentration of 9,5 μM RNP complex concentration (molar ratio 2:1, 19 μM sgRNA mix: 9,5 μM Cas9) and incubated at 37 °C for 5 min then cooled on ice and stored in 4 °C until microinjection. Approximately 1 nL of RNP complex composed of three sgRNA targeting different gene loci and Cas9 protein was injected into the yolk of one-cell stage embryos [59]. Uninjected siblings were used as control. In the case of both *atp1b3b* and *colec10* knockdown, injection of 28.5 µM of Cas9-sgRNA complex according to previous report [57] induced over 70% mortality which suggests batch-sepcific toxicity of the Cas9 enzyme (not shown), therefore we used 9.5 µM in our experiments. To facilitate the observation of the heart, all experiments were performed in the transgenic line *Tg(myl7:EGFP)* [39] or *Tg(myl7:EGFP-Hsa.HRAS)^s883^* [40] expressing EGFP in the heart. To account for possible off-target effects resulting from microinjections, we targeted *EGFP* in the same transgenic zebrafish lines using *in vitro* transcribed sgRNAs [59]. Templates for sgRNA synthesis were generated by annealing and elongation (Phusion Flash High-Fidelity Master Mix, Life Techologies) of a forward primer containing a T7 promoter and guide sequence, and a reverse primer encoding the sgRNA scaffold (Supplementary Table 7). The DNA templates were purified (MinElute PCR Purification, Qiagen) and quality checked by 8% polyacrylamide gel electrophoresis before used as template for *in vitro* transcription using T7-FlashScribe™ Transcription Kit (CellScript). Obtained sgRNA mix was purified using RNA Clean and Concentrator-5 (Zymo Research). Injections were performed as before with the exception that 60 μM sgRNA mix was combined with diluted 10 μM Cas9 protein to reach final ∼5 μM RNP complex concentration (molar ratio 6:1, ∼60 μM sgRNA mix: 10 μM Cas9).

### Knockout phenotype and heart rate analysis

Following microinjection, dead embryos before 24 hpf were discarded to exclude unfertilized embryos or those damaged by the injection. Phenotypic analysis was performed starting from 48 hpf. First, malformed embryos and/or embryos that developed cardiac edema were separated and reported. Then, wherever possible, a random 10 and 20 embryos from the uninjected control and knockout groups were sampled for heart rate analysis and kept for further observations. Heart rate analysis was performed at 48 hpf on the SZX16 fluorescent steromicroscope (Olympus) by manually counting the number of heartbeats for 30 seconds per embryo using a timer. Subsequent statistical analysis was performed in R. Shapiro-Wilk normality test implied that the distribution of the knockout data are significantly different from a normal distribution (p-value < 0.05), therefore, non-parametric Wilcoxon rank sum test was applied to compare data between wildtype and knockout embryos. The Holm-Bonferroni method was used for adjusting p-values. All movies were recorded using acA1440-220um USB 3.0 camera (Basler).

## Results

### Isolation and transcriptome profiling of single cells from the embryonic zebrafish heart

To generate homogenous, viable single cell suspension from the zebrafish heart at 48 hpf and 72 hpf, we optimized a cell dissociation protocol incorporating simultaneous trypsin and collagenase treatment. We obtained cell suspension with viability above 90% (Supplementary Fig. 1A). To provide an internal control of specific rare cardiac cell population, we utilized two transgenic lines *sqet33mi59B* [32,37] and *sqet31Et*, in which cells of the sinoatrial (SA) and atrioventricular (AV) pacemaker regions expressed EGFP [33,36]. In order to additionally demarcate the major cell types of the heart, we utilized the *Tg(myl7:mRFP)* transgenic line [38] to confidently mark cardiomyocytes (CMs), which is the most technically challenging cell type to isolate, and enhance cell clustering. To profile the transcriptome of the zebrafish heart, we collected offspring from two independent crosses of *Tg(myl7:mRPF)* line with either *sqet33mi59BEt* or *sqet31Et* transgenic lines. From each of the double transgenic individual pools, we isolated whole hearts at 48 hpf and 72 hpf and dissociated them into single cells which were subsequently encapsulated according to the 10x Genomics workflow (Fig. 1A).

In total, we obtained 52 447 cells from all four biological samples (Fig.1B, Supplementary Table 1). In each, over 22 000 genes were identified. The median number of genes per cell varied between the developmental stages, which was on average nearly three times higher at 48 hpf as compared to 72 hpf. Similar trend was observed in terms of unique transcript molecules per cell, whereas mitochondrial gene content was double in cells derived from 72 hpf (Supplementary Fig. 1B). At the same time, the Pearson correlation coefficient between raw developmental pseudo-replicates was equal to 0.98 and 0.83 for 48 hpf and 72 hpf data, respectively (Fig.1D). The number of cells after quality control for library size, mitochondrial gene content and possible multiplets ranged from 6728 to 11 015 per sample, resulting in a final number of 34 676 cells for downstream analysis (Fig.1B, Supplementary Table 1). Out of these, 2243 (80% of total *RFP*+ cells) cells co-expressed *RFP* and well established gene signatures for myocardial cells which included *cmlc1* and *myl7*. These form a separate cluster, suggesting cardiomyocytes identity (Fig. 1C, Supplementary Fig.1C). In addition, 179 cells were found to express *EGFP,* out of which the vast majority comes from 48 hpf dataset (129 cells at 48 hpf vs. 50 cells at 72 hpf). Most of the total EGFP+ cells (116) were within the cardiomyocytes cluster, suggesting that our protocol enabled the isolation of single cardiomyocytes, with sufficient sensitivity to capture rare cell populations making up the two pacemaker regions. To facilitate in-depth exploration of our data, we developed an R Shiny application based tool for interactive data visualization, differential expression as well as gene enrichment analysis which is available at http://zfcardioscape.iimcb.gov.pl.

### The cellular landscape of the developing zebrafish heart

We distinguished 18 discrete cell populations comprising the zebrafish embryonic heart from both 48 hpf and 72 hpf (Fig. 1C). The most abundant clusters comprised endothelial cells (31%), cardiomyocytes (11%) and mesenchymal fibroblasts (7%) (Fig. 1E, Supplementary Table 2). Moreover, the number of cells contributing to each cluster varied between the developmental stages. Clusters originating mostly from 48 hpf included resident fibroblasts, mesoderm progenitors and cardiac peripheral nerves. On the other hand, seven clusters showed a higher proportion of cells derived from 72 hpf. These included neuronal cells, epicardial cells and Bulbus arteriosus (Fig. 1E, Supplementary Table 2). Of note, mitochondrial gene content was particularly higher among cells originating from the epicardium and cardiomyoytes clusters in the later developmental stage (72 hpf) which suggested their higher energy metabolism. On the other hand, the number of expressed genes and UMIs were greater at 48 hpf as compared to 72 hpf. Among clusters with the highest number of gene and UMIs were Mesoderm progenitors, AV endocardium and Neural crest cells (Fig. 1F).

Based on the expression of unique marker genes, these clusters could be broadly grouped into the major cell lineages of the developing heart, namely: myocardial, endocardial, epicardial, neural and neural crest, and mesenchymal/fibroblasts. The myocardial lineage formed a distinct cluster of cells represented by the cluster “Myocardium” which could be distinguished by the expression of *myl7, cmlc1, myh6, myh7, tnnt2* (Fig.2A, Supplementary Table 3) [60,61], as well as the *RFP* transgenic marker (Supplementary Fig. 1C). The endocardial lineage constituted the largest proportion of the cells in our data. These included the clusters “Endothelial cells” and “Endothelial precursors” which express *cdh5*, *tie1* and *fli1* [62,63], as well as the “AV endocardium” and “AV cushion” which contributes to the developing AV valve (Fig. 1E, Supplementary Table 2). In addition to endocardial markers, the latter two express AV canal restricted markers such as *anxa5b*, *alcama*, and *nrg1* [31,33]. The “AV cushion” differs from the “AV endocardium” cluster by the presence of myocardial-associated genes (*myl7*, *ttn.2*, and a number of mitochondrial genes) and a higher expression of cell adhesion and extracellular matrix proteins (*postna*, *col1a1a/b/2*) (Supplementary Table 3). The epicardial lineage forms a single cluster “Epicardium” characterized by the expression of *tbx18, gstm.3, wt1b,* and *tcf21* [28].

**Figure 2.**
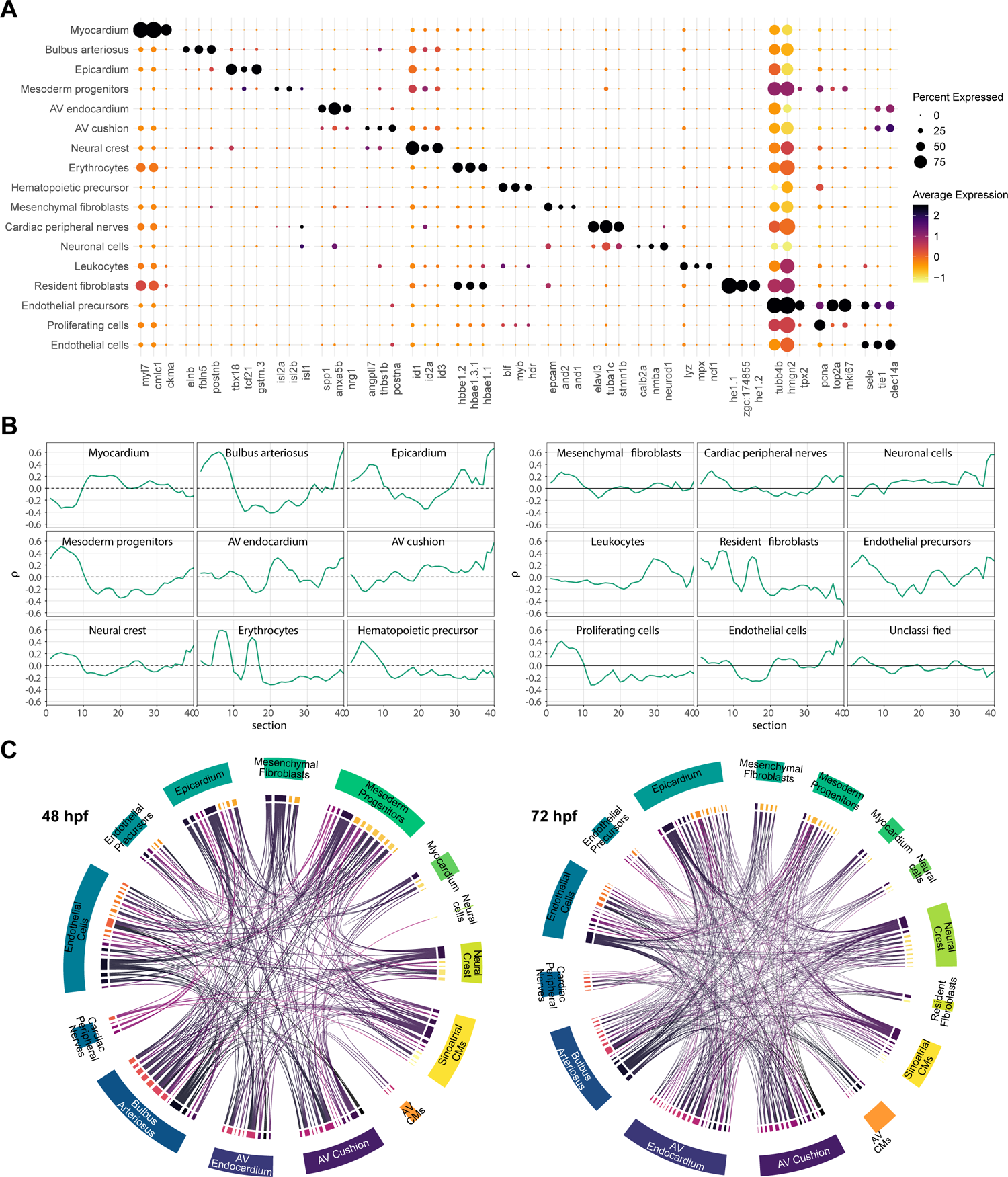
The expression profiles of main cardiac cell clusters correlate with heart structures of known spatial localization. **A** Dotplot showing the top three differentially expressed genes for all cell clusters except “Unclassified”. **B** Spatial projection of cardiac single cell expression profiles to zebrafish heart sections derived from Burkhard et al. [31] at 48 hpf. **C** Interaction map between cell clusters according to ligand-receptor expression. Each line connect individual ligand with its corresponding receptor (source data: Supplementary Table 4).

Cells of the “Bulbus arteriosus” cluster were distinguished by the expression of *elnb, postnb,* and *fbln5*, reflecting their smooth muscle composition [19]. Neural and neural crest cells were represented by clusters “Cardiac peripheral nerves”’ and “Neuronal cells”’ characterized by the expression of *tuba1c* and *elavl3* [64] and cluster “Neural crest” by the expression of *twist1a/b, id2a, id3*, and *sox9b* [65–67]. Mesenchyme and fibroblasts make up a significant population of cardiac cells and could be distinguished by the expression of adhesion molecules such as *cldnh,* and *epcam* [68]. Two clusters constituting this cell lineage were “Resident fibroblasts”and “Mesenchymal fibroblasts”, the latter of which expressed higher numbers of genes encoding structural proteins and extracellular matrix components such as *col1a1a*, *cdh1,* and those encoding the Claudin family [69] (Supplementary Table 3). Interestingly, we also detected a cluster of cells designated as “Mesoderm progenitos” (Fig. 1C, 2A) which was enriched in the expression of *isl1* and *isl2b*, and *ldb1a* which are known to be expressed in cells originating from the second heart field [16,70]. Cells of this cluster also expressed a large number of transcription factors and cell cycle regulators (Supplementary Fig. 1D, Supplementary table 3). Likely as byproduct of our whole heart isolation procedure, we also captured hematopoietic cells indicated by the expression of *blf* and *myb* [71,72]. From these, distinct population of erythrocytes (expressing hemoglobin genes as well as *alas2, hemgn* [73]) and leukocytes comprising of neutrophils (*mpx* [74], *ncf1* [75]), macrophages (*mpeg* [76]), and platelets (*mpl, itga2b* [77]) could be distinguished based on the expression of their unique markers (Fig. 2A, Supplementary Table 3).

Different cell clusters exhibited variable proliferation activity based on the expression of genes driving the G2M and S-phase of the cell cycle. The most proliferative cluster being “Endothelial precursors”, the cells of which expressed a large number of G2M and S-phase related genes (Supplementary Fig. 1D). The clusters “Bulbus arteriosus”, “Leukocytes”, “Mesoderm progenitors’’, “Erythrocytes”, and “Resident fibroblasts” were found to be more proliferative at 48 hpf compared to 72 hpf (Supplementary Fig. 1D), which may reflect their differentiation process. Accordingly, clusters expressing known cell-type-specific markers were generally low in proliferation activity (<50%). These include “Myocardium”, “AV cushion” and “AV endocardium”, “Endothelial cells”, and “Neuronal cells” (Supplementary Fig. 1D).

In order to provide spatial context to the cell clusters, we took advantage of the available spatial information from the serial cross-section transcriptome dataset of the zebrafish heart [31]. Correlating the expression profiles of our main scRNA-seq clusters with that of each serial section, the analysis revealed that cell types associated with structures known to be spatially localized exhibited the highest correlation with sections representing their corresponding localization (Fig. 2B). Furthermore, spatial correlation between each main cluster and their corresponding fine-grained clusters (see Supplementary method) revealed overall high levels of both gene and spatial cluster correlation, except for the “Unclassified” cluster which was heterogenous in both spatial and gene expression values, and the “Mesenchymal fibroblasts” cluster which was homogeneous in terms of gene expression values but spatially heterogeneous (Supplementary Fig. 2, Supplementary Table 6). This suggested that these clusters might be controlling subparts of the main clusters which have a differentiated role but are otherwise attached to the developmental structure associated with the respective main cluster. Taken together, the spatial correlation analyses of the scRNA-seq clusters agrees with the spatially restricted expression profile, thereby providing additional support for our cluster annotation.

To identify potential cellular crosstalk between cell types in the developing heart, we constructed an interactome map between the cardiac cell clusters based on the expression of ligands and their receptors [55]. The analysis revealed several known cellular interactions including those involved in the development of AV cushion [78]. Genes encoding Bmp4/5/6 were expressed in the “AV cardiomyocytes” cluster, while those encoding its signaling receptors Bmpr2a, Acvr1ba, and Sdc2 were expressed in the “AV cushion” cluster (Fig.2C, Supplementary Table 4). The analyses also suggested potential interactions which were not previously established. For instance, the cluster “Cardiac peripheral nerves” interacted with “Endothelial cells” and “Epicardium” clusters through Robo1/2 and its receptor Slit1a/2/3 expressed in the corresponding clusters (Fig. 2C, Supplementary Table 4). The Robo/Slit signaling was implicated in axon guidance, as well as in endocardial progenitors migration to the midline [79]. The “Cardiac peripheral nerves” cluster also interacted with “SA cardiomyocytes” subcluster through several known ligand-receptor pairs including Plxna3 - Sema3aa, Fgfr2/3/4 - Ncam1a, and Sdc4 - Mdkb (Fig.2C, Supplementary Table 4), which have been implicated in neural patterning [80,81]. Taken together, the analysis revealed potential interactions between cell or tissue types and their underlying molecular players for further in-depth analyses.

### Diverse myocardial cell populations constitute chamber myocardium and the cardiac conduction system

The myocardium is the main tissue type of the heart which is responsible for the organ’s main contractile function. Myocardial cells are characterized by their contractility and high metabolism [82]. In agreement with this, Gene Ontology terms related to muscle function and structure were enriched among genes in this cluster (Figure 3A, B). Chamber cardiomyocytes could be distinguished by the expression of well-established marker genes, namely atrial *myh6,* and ventricular *myh7/myh7l* [83,84]. Within the cluster “Myocardium”, a clear distinction between atrial and ventricular cardiomyocytes was evident by the complementary expression of atrial (*myh6*) and ventricular (*myh7* and *myh7l*) cardiomyocyte markers (Fig. 3A). Differential expression analysis between both fractions confirmed the presence of a set of unique genes in each heart chambers. The top differentially expressed signatures comprised genes encoding for various structural proteins involved in active force generation, including *myh6, tnnc1b* and *smtln1* (atrial), and *myh7l*, *myh7*, and *tnni4a* (ventricular) (Fig. 3C).

**Figure 3.**
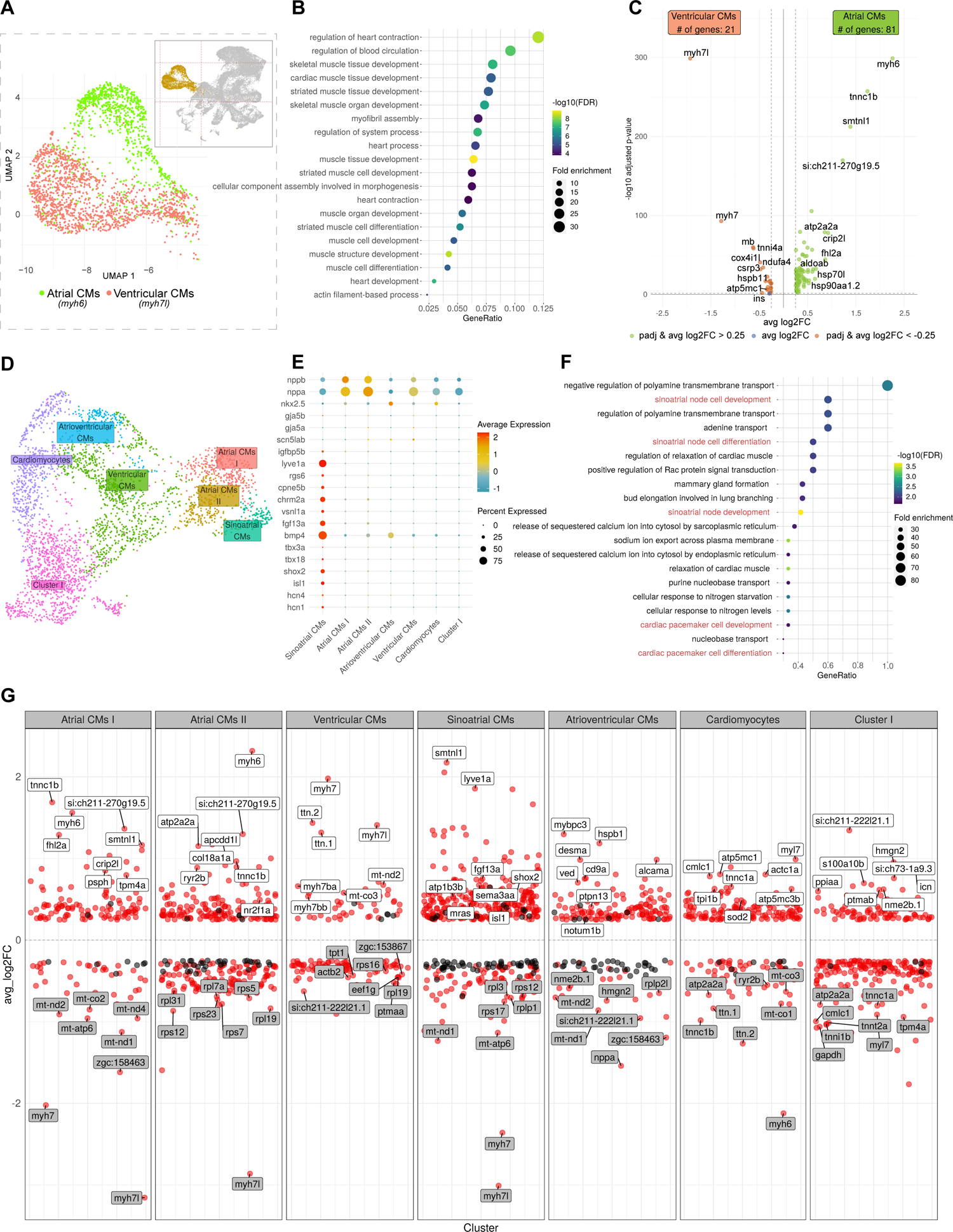
Analysis of the “Myocardium” cluster revealed the diversity of myocardial cells and pinpointed sinoatrial cardiomyocytes within the atrial myocardium. **A** Two distinct identities of chamber myocardium could be delineated by molecular markers: Atrial CMs by expression of *myh6* and Ventricular CMs by *myh7l* expression. **B** Gene Ontology enrichment analysis of all genes enriched within the “Myocardium” cluster. **C** Volcano plot depicting differentially expressed genes between atrial and ventricular CMs fractions. **D** UMAP projection of re-clustered myocardial cells reflecting the heterogeneity of atrial and ventricular myocardium. **E** Dotplot showing the expression of well-established gene signatures associated with working myocardium and sinoatrial pacemaker within “Myocardium” clusters. **F** Gene Ontology enrichment analysis of genes enriched within the “Sinoatrial CMs” subcluster. **G** Differentially expressed genes in each myocardial subclusters. Top 8 genes in terms of significance were labeled. Genes with adjusted p-value < 0.05 are depicted as red dots while not significant genes (adjusted p-value > 0.05) are shown in black. Source data: Supplementary Table 10, 11.

**Figure 4.**
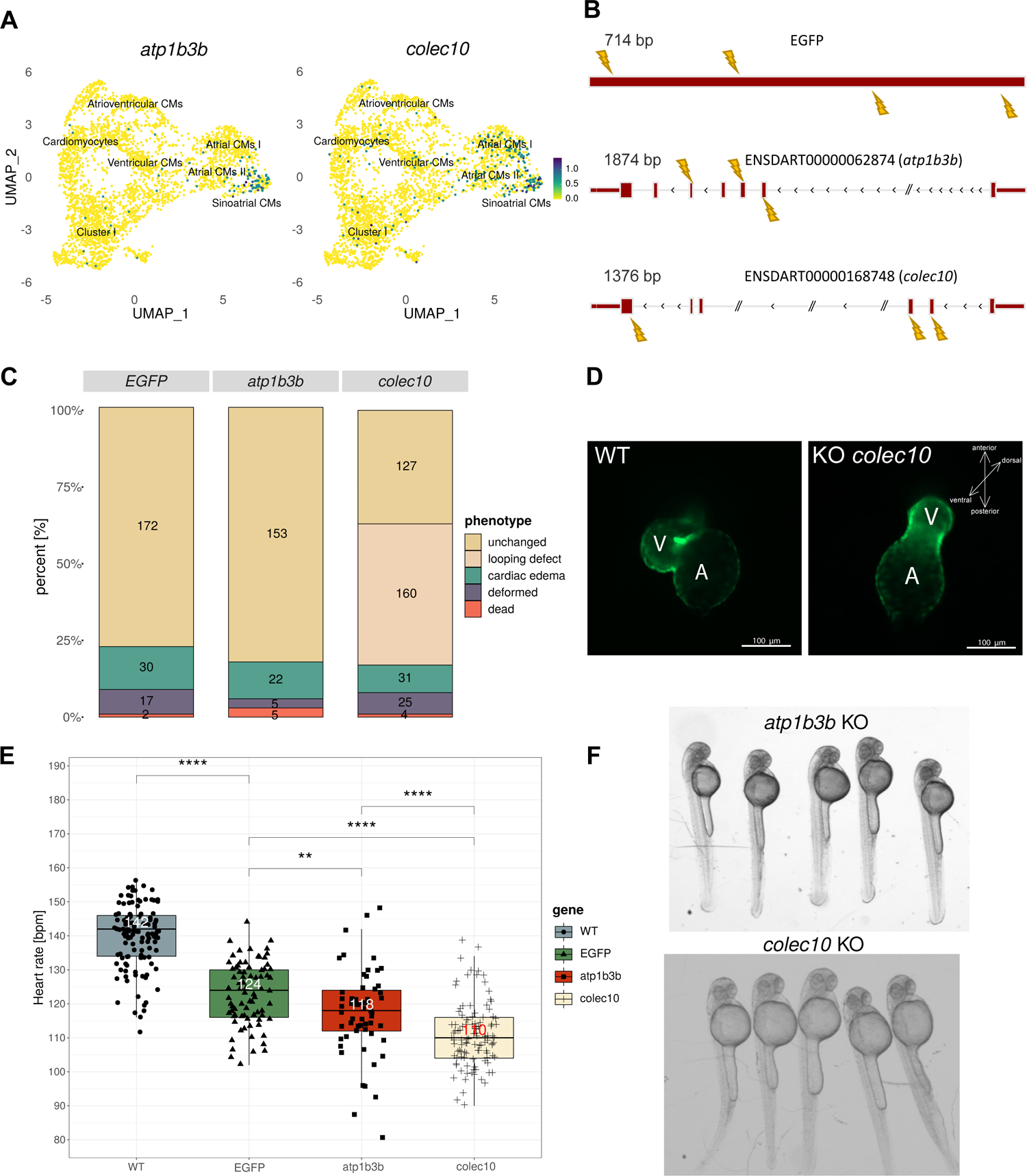
Functional analysis identified atp1b3b and colec10 as new players in heartbeat maintenance. **A** Expression of *atp1b3b* and *colec10* within the myocardial subclusters. The highest expression for both genes was observed within Sinoatrial CMs cluster. **B** Scheme illustrating gene loci targeted by CRISPR/Cas9 strategy. The combination of four sgRNAs was used to target EGFP whereas three sgRNAs were utilized for *atp1b3b* and *colec10*. **C** Barplot summarizing observed phenotypes from all replicates for each gene knockout. Numbers within each section indicate the number of embryos associated with a particular phenotype. **D** Fluorescent images showing heart looping defect in *colec10* knockouts as compared to uninjected embryos (WT represents embryos on *Tg(myl7:EGFP-Hsa.HRAS)^s883^* background at 48 hpf). **E** Heartbeat rate comparison between appropriate knockouts, uninjected wild-type control, and EGFP-injected embryos (significance values according to Wilcoxon rank sum test). **F** Representative image showing morphology of *atp1b3b* and *colec10* knockout embryos.

Further re-clustering of the cells within the “Myocardium” cluster (a total of 3315 cells) revealed subpopulations of myocardial cell types (Fig. 3D). At least 43% of myocardial cell populations consisted of those making up the working myocardium of the two cardiac chambers, namely, atrial (“Atrial CMs I” and “Atrial CMs II”) and ventricular (“Ventricular CMs”). The mutually exclusive expression of *myh6* and *myh7/myh7l*, together with other unique markers, differentiate these two subtypes of chamber myocardium (Fig. 3G, Supplementary Table 5). Two distinct subclusters of atrial cardiomyocytes were different in terms of higher mitochondrial content in “Atrial CMs II” compared to “Atrial CMs I” (Supplementary Fig. 1E). In addition, “Atrial CMs II” was enriched for the expression of *myoz2b*; Supplementary Table 5), the mammalian ortholog of which was previously reported to mark a distinct population of cardiomyocytes with yet unknown function [85]. An additional cluster designated as “Cardiomyocytes” expressed *myl7* which identified these cells as cardiomyocytes. However, they are not enriched in any chamber-specific markers.

Besides the working myocardium, the cardiac conduction system constitutes an important structure of myocardial origin. Among cells with high expression of atrial-specific gene (*myh6*), we identified a group of 127 cells that uniquely express *lyve1a* [31]*, vsnl1a* [86], and *chrm2a* [87] that have been reported to be upregulated in the heart SA region. These cells also express other genes implicated in the function of the SA pacemaker, including *isl1, shox2, hcn4, bmp4, fgf13a, tbx18* [31,32,88], while maintaining low expression of working myocytes markers, e.g. *nppa/b, nkx2.5, gja5a/b* (Fig. 3E). Gene Ontology enrichment analysis revealed numerous GO terms related to the sinoatrial node and cardiac pacemaker cell development (Fig. 3F). We therefore define this cluster as “Sinoatrial CMs” that could represent a group of specialized cardiomyocytes of the primary pacemaker site of the cardiac conduction system. Similarly, the cluster “Atrioventricular CMs” is distinguished by the expression of *alcama*, *bmp4*, and *tbx2b* (Fig. 3E, G, Supplementary Table 5), which were previously associated with the atrioventricular canal [33].

### Functional analysis of cell-type-specific markers reveal new potential players in heart rhythm maintenance

Single cell analysis allowed us the opportunity to identify and study molecular components implicated in the development and function of rare cell populations. To this end, we utilized the CRISPR/Cas9 based knockdown [57,59] to target two candidate genes, *atp1b3b* and *colec10*, which were significantly enriched in the “Sinoatrial CMs” cluster representing the rare population of primary pacemaker cells (Fig. 4A, Supplementary Table 5). To target *atp1b3b,* we designed sgRNAs against the 2nd, 4th and 5th exons (Fig. 4B). By 48hpf, loss of function of *atp1b3b* did not appear to affect heart development (Fig. 4D). Moreover, these embryos also looked largely unaffected in terms of overall morphology (Fig. 4C, F). Targeting of *colec10* by sgRNAs directed against the 2nd, 3rd and 6th exons caused a high proportion of embryos with hearts that failed to loop by 48 hpf (46 %, n = 160/347, Fig. 4C). These embryos appeared morphologically normal otherwise, with only a very small number of embryos exhibiting gross morphological abnormality (Fig. 4C, F).

The specific expression of *atp1b3b* and *colec10* in the “Sinoatrial CMs” representing the primary pacemaker cell population led us to question whether they are involved in regulating heart rhythm. We systematically calculated the heart beat rate in CRISPR/Cas9-injected embryos as compared to uninjected siblings and EGFP-knockout embryos (EGFP-KO) which served as controls (Figure 4E, Supplementary Fig. 3A). Injection of Cas9 complex with sgRNAs targeting *EGFP* successfully abolished 100% (n = 77) EGFP expression in the transgenic embryos. A small number developed cardiac edema (13,5%, n = 30/221) while the majority did not show any observable phenotypic changes (Fig. 4C). Interestingly, we observed a reduction in heart rate in EGFP-injected embryos as compared to uninjected siblings (Supplementary Fig. 3A). Therefore, to account for off-target effects resulting from microinjections, in addition to uninjected siblings, we directly compared heart rates of *atp1b3b* and *colec10* knockout embryos to EGFP-injected ones. We observed that, in both cases, heart beat rate was significantly lower by 48 hpf (Fig. 4E, Supplementary Movie 1-2). Knockout of *atp1b3b* resulted in a median of 118 bpm which was 24 and 6 heart beats per minute (bpm) less compared to uninjected siblings (n = 61, p-value = 2.2e-16) and EGFP-KO embryos (p-value = 0.005543); whereas for *colec10* knockout, the larvae had 110 bpm which is 32 and 14 bpm less compared to uninjected siblings (n = 109, p-value = 2.2e-16) and EGFP-KO embryos (p-value = 5.419e-14) (Fig. 4E, Supplementary Fig. 3A). Moreover, some of the *atp1b3b* knockout embryos developed arrhythmia later on (by 72 hpf or 96 hpf) characterized by either irregular heart rhythm or silent ventricle (at least 4 out of 50 embryos observed; Supplementary Movie 3-4). The reduced heart rate in EGFP knockouts could possibly result from developmental delay triggered by microinjection. Nevertheless, direct comparison between embryos injected with sgRNAs targeting either *atp1b3b* or *colec10* and *EGFP* still indicated significantly lower heart rate in the former (Fig 4E), thus suggesting the specificity of heart rate phenotype to the knockdown of both genes. The genetic mozaicism of these F0 individuals limited our ability to ascertain the true function of the two candidate genes. Hence, more detailed analysis of stable mutants is necessary and underway. Collectively, these observations suggest that the two newly identified sinoatrial pacemaker genes play a role in heart development and regulation of heart rhythm.

### The cardiac endothelial cell subpopulations possess distinct molecular profiles

Endothelial cells perform critical function in heart development and physiology and are known to contribute to different heart tissues including cardiac valve, coronary vessels, and trabeculae [89]. In order to characterize the diversity of cardiac endothelial cells in the developing heart, we re-clustered cells of the “Endothelial cells”, “Endothelial precursors”, “AV endocardium”, and “AV cushion” clusters. Our analyses revealed that the cardiac endothelial cells consisted of molecularly distinct populations (Fig. 5A). Clusters “AV endocardium” and “AV cushion” were distinguished by the expression of several genes characteristic of the endocardium of the AV region, which included *hand2, has2, alcama,* and *crip2* [33,78,90] (Fig. 5B). Cluster “AV endocardium” genes included *anxa1a* and *anxa5b*, both of which are known to be expressed in AV canal region [31], as well as *notch1b* and *klf2a* which are early markers of the AV valve [78] (Fig. 5B). Cluster “AV cushion” was enriched in genes coding for extracellular matrix constituents including many collagen family (*col1a1b, col1a2*, *col5a1/2a,* and *thbs1a/b*), as well as *postna* which are known to be expressed in the AV cushion [33]. This cluster also had the highest expression of *col1a2* which is implicated in human cardiac valve disease [91] (Fig. 5B, Supplementary Table 8).

**Figure 5.**
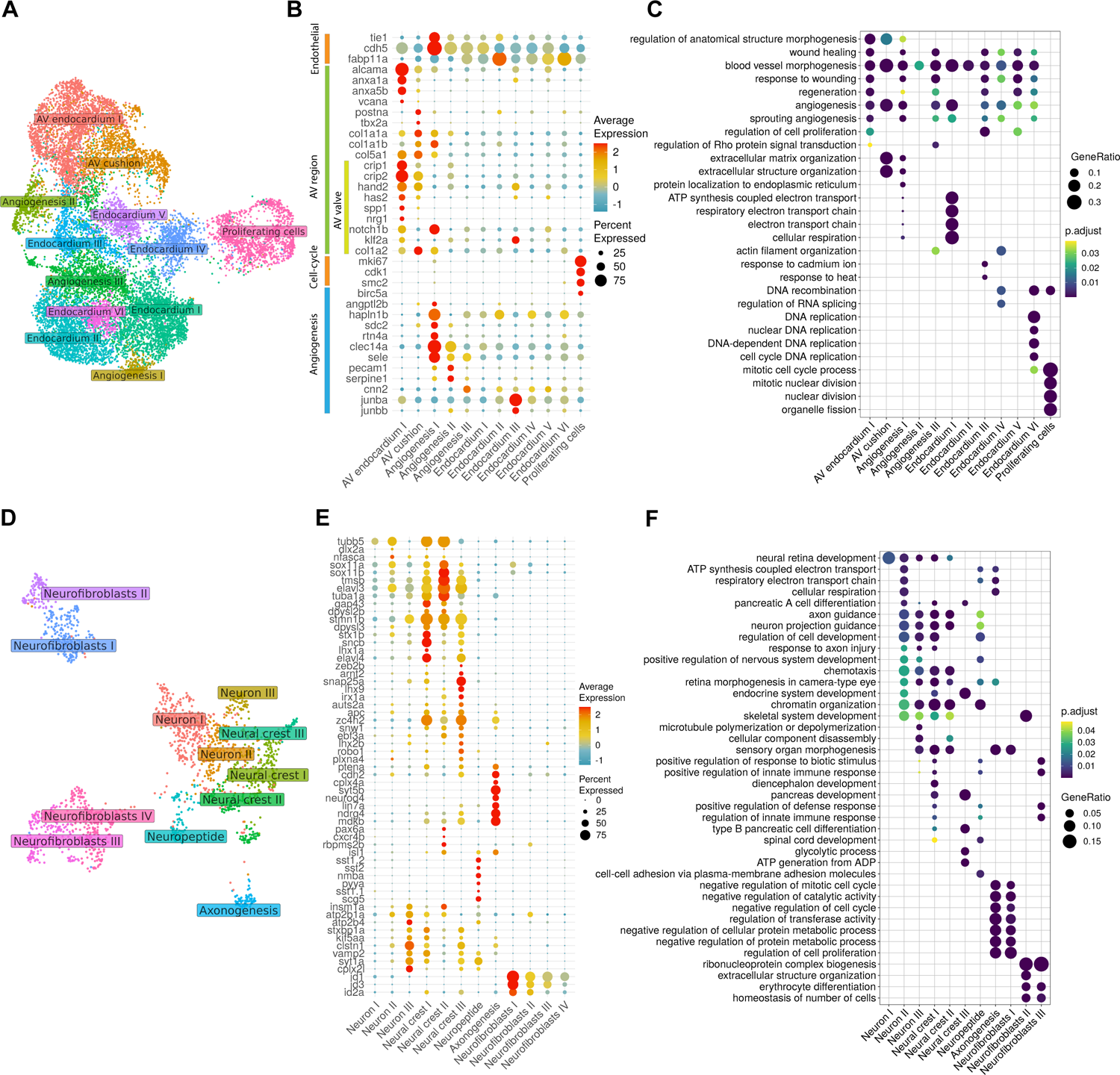
Molecular profiles of diverse endothelial and neural cell subclusters. UMAP projection of endothelial **(A)** and neural **(D)** subclusters. Dotplot showing the expression of specific marker gene signatures associated with various endothelial **(B)** or neural **(E)** cell subclusters. Average expression values represent normalized counts, percent expressed values represent the proportion of cells within a cluster that expresses a particular gene. Gene Ontology enrichment analysis of each endothelial **(C)** and neural **(F)** subclusters depicting the enriched functional terms (GO level 5) in each cluster. Adjusted p-values (Holm–Bonferroni) are indicated in colour, gene ratio represents the gene count mapped into each GO category. Source data: Supplementary Table 12, 13.

Noticeably, several of the endothelial subclusters were significantly enriched in functional terms related to blood vessel morphogenesis and angiogenesis (Fig. 5C, Supplementary Table 12). Cluster “Proliferating cells” was significantly enriched in the expression of genes encoding cell cycle regulators (Fig. 5C, Supplementary Table 8). Cells of this cluster uniquely express *birc5a* which is known for its role in vasculogenesis and angiogenesis [92]. Cells of “Angiogenesis I” cluster expressed the most number of genes implicated in sprouting angiogenesis, including *hapln1b* [93]*, sdc2* [94], and *rtn4a* [95] (Fig. 5B). Cluster “Angiogenesis II” shared a partially overlapping set of angiogenesis genes with “Angiogenesis I”, such as *clec14a* [96], as well as *sele* [97], *pecam1* [98] and *serpine1* [99]. Apart from *sele* and *serpine1*, cluster “Angiogenesis III” also expressed several unique genes including *cnn2* which is implicated in blood vessel endothelial cell migration [100] (Fig. 5B). The rest of the endothelial cell clusters were not enriched in any specific functional category other than that which suggest their endothelial identity and hence are likely to be endocardial lining of the cardiac chambers. Collectively, the diversity of the cardiac endothelial cell population reflects their contribution to various structures of the heart and suggests their functional diversity.

### Neural crest and the intracardiac nervous system

Besides the main cardiac cell lineages, we also identified other cell populations whose expression profiles we provide as a resource (Supplementary Table 9). One important but poorly characterized cell populations are those making up the cardiac peripheral nervous system, which ensures the regulation of heart contraction speed according to physiological demands [101]. To evaluate the diversity of the neural and neural crest cell populations in the zebrafish heart, we re-analyzed the main clusters “Cardiac peripheral nerves”, “Neural crest” and “Neuronal cells”. We distinguished subpopulations of cells which could be broadly subdivided into neuronal and non-neuronal cells (Fig. 5D). Clusters “Neurons I-III” and “Neural crest I-III” were enriched in the expression of neurodifferentiation markers including *elavl3, elavl4, stmn1b, tuba1a,* and *tubb5* [64,102–104] (Fig. 5E, F). They were also enriched in the expression of *sncb* which is involved in dopaminergic neuron differentiation and development of motor functions [105]. Clusters “Neural crest I-III” were additionally enriched in the expression of several genes implicated in neural crest development, including *mdkb*, *pbx4*, *tfap2a*, and *snw1* [106–109]. Cluster “Neuropeptide” was enriched in the expression of genes encoding neuropeptide signaling molecules (*npb* [110]*, scg5* [111]*, nmba,* and *pyya* [112]), as well as factors promoting dendrite morphogenesis and cell migration (*sst1.1/1.2/2* [113], and *tmsb* [114]) (Fig. 5E, Supplementary Table 9). The “Axonogenesis” cluster expressed genes encoding proteins known to be associated with axons or denrites including *dscaml1, tnc, draxin,* and *nptnb* [115–117].

Compared to other neural subclusters, clusters “Neurofibroblasts I-IV” were depleted in neural and neural crest markers expression while enriched in the expression of *id1* and *id3* known for their role as inhibitors of neurogenesis [118] (Fig. 5E, Supplementary Table 9). Clusters “Neurofibroblasts I” and “Neurofibroblasts II” expressed a large number of genes implicated in cell cycle such as *jun/ba/bb/d* and *cdkn1bb*. They also uniquely expressed *egr1* which is implicated in neural stem cell maintenance [119], suggesting the characteristics of neuron stem-like cells. Hence, their expression profiles suggest their identity as neural fibroblast cells. Cluster “Neurofibroblasts III” was enriched in genes coding for the most number of extracellular matrix components including collagens and keratins, which suggests structural function. Together with “Neurofibroblasts IV” cluster, it was also enriched for *postnb* encoding the zebrafish ortholog of Periostin which is known to be expressed in endoneurial cardiac fibroblasts contributing to sympathetic nerve fasciculation in the mammalian heart [120]. Taken together, the expression resource for these clusters, as well as others, could potentially reveal novel cardiac cell types which are poorly explored and open new pathways of investigations.

## Discussion

We present a high resolution atlas of the developing zebrafish heart between 48 hpf and 72 hpf obtained through single cell transcriptomics. The developmental stages chosen constitute a critical phase of heart development, where the development of specialized cardiac structures occur, including the formation of atrioventricular canal, heart looping, as well as the development of the cardiac valves and conduction system [78,121]. In addition, this developmental period also covered the time when external cell types are incorporated into specialized structures of the heart, such as the neural crest and the proepicardium [7,17]. Our study delineated major cardiac cell lineages which were supported by known molecular markers. In addition, spatial correlation analysis points to main clusters which possess specific anterior-posterior localization within the heart tube, which provided further support of their identity. Further correlation analysis between the main and subcluster components (fine-grained clusters at higher resolution, see Supplementary Methods) revealed the homogenous composition of several main clusters, while pointing out the heterogenous composition of other clusters, such as the mesenchymal fibroblasts, on which a scarce literature information is available and annotation is therefore less precise. To date, there is only one published spatial transcriptomic data on the embryonic zebrafish heart [31]. Although the section-based method limits the cellular resolution, it still provided a reliable positional reference for major cardiac structures. Future spatial analysis at higher resolution is expected to enable the annotation for each cell type with improved accuracy.

Our analyses uncovered subpopulations within the major cardiac cell lineages, including previously uncharacterized subtypes of cells, along with their molecular profiles. Previous single cell analyses on the developing epicardium in zebrafish has uncovered distinct subpopulations, each possessing specific genetic signatures and function [28]. Similarly, transcriptome profiling of mammalian cardiomyocytes at the single cell level have revealed unprecedented molecular diversity which seems to suggest further functional specializations and/or distinct energetic profiles within this cell type [25,85,122]. A specific population of cardiomyocytes that express Myoz2, a member of the sarcomeric protein family that is implicated in hypertrophic response [123], was recently described in mouse and humans [26,85]. We observed that *myoz2b,* the zebrafish ortholog of this gene was expressed in the zebrafish myocardium. *myoz2b* expression were detected in subsets of cells across almost all myocardial clusters. Determining the features and function of this particular cardiomyocyte population would provide key insights into how it contributes to the overall functioning of the heart.

A similar functional heterogeneity was also observed among cardiac endothelial cells. An interesting observation is that several endothelial cell clusters were found to be enriched for genes associated with blood vessel formation and angiogenesis. It was previously reported that the zebrafish coronary vasculature originates from the endocardial population and develops only from 1-2 months post-hatching [124]. Moreover, endocardial cells are known to share the expression of vascular markers [125]. Therefore it would seem unlikely that the enrichment of these functional categories reflects an actual process of vasculogenesis. Nevertheless, the distinct molecular profiles of the cardiac endothelial cell populations suggest the presence of functional diversity at the embryonic stage of heart development.

Our single cell transcriptome analyses also captured diverse populations of at least two extracardiac cell lineages - the cardiac neural crest and fibroblasts. Annotation of these cell types, particularly the latter, are still challenging as there are currently very few molecular markers that can be used to define them [8]. Previous studies in the zebrafish have focused on cardiac fibroblasts within the context of regeneration and hence capture only the activated form [126,127]. We nevertheless provide their expression profiles as a resource which is envisaged to open the pathway for further studies of various aspects of cardiac biology in the zebrafish model organism.

The upcoming challenge will be to ascribe function to each cell type and determine the extent to which these diversity of cell types are conserved in comparison to the human heart in terms of their molecular properties and function. As a an initial step towards this effort, we functionally characterized two genes, *atp1b3b* and *colec10*, which were specifically enriched in the “Sinoatrial CMs” cluster representing the primary pacemaker region. Our loss-of-function analyses revealed the potential role of both genes in heart development and maintaining heart rhythm. The *atp1b3b* gene encodes the ꞵ3 subunit of the Na+/K+ ATPase family responsible for osmoregulation and electrical excitability of nerve and muscle. In the heart, the ɑ1 and ɑ2 isoforms, as well as the ꞵ1 are predominantly expressed [128]. However, some studies have also reported the expression of ꞵ3 isoform in the ventricular myocytes [129]. Evidence have suggested that a functional distinction exist between the isoforms, although the exact mechanism is still poorly understood. Interestingly, our single cell transcriptome analysis showed that *atp1b3b* had the most restricted expression in the sinoatrial CMs compared to other genes encoding the ɑ and ꞵ subunits. This may also underlie the specific heart rhythm phenotype observed upon their loss of function. Future analysis will further distinguish the function and mechanism of the different members of the Na+/K+ ATPases which has potential implications in developing therapies for heart failure. The gene *collectin sub-family member 10 (colec10)* encodes a member of the C-type lectin family [34], mutations of which have been associated with the rare 3MC syndrome 3 in humans [130]. The gene product of *colec10* was implicated in cellular movement and migration in vitro [130], while scarce information is available on the function of this gene in vivo. Our observation of its specific expression in the sinoatrial CMs, together with the defects observed upon its loss of function, therefore suggest that this gene may be a new player in regulating heart development and/or function.

## Acknowledgements

We are grateful to the zebrafish core facility of the IIMCB Warsaw for excellent fish care; V. Korzh, A. Minoda, T. Kelly, R. Foo, as well as all members of the C.W. lab especially M. Łapiński for fruitful discussions and critical guidelines. The project POIR.04.04.00-00-1AF0/16-00 carried out within the First TEAM program of the Foundation for Polish Science co-financed by the European Union under the European Regional Development Fund support C.W., K.A.N., and A.S.. R.H. and M. Z. were supported by a grant from the Priority Research Area DigiWorld under the Strategic Programme Excellence Initiative at Jagiellonian University. M. Z. was supported by the Polish National Agency for Academic Exchange and by National Science Center Poland (2021/42/E/NZ2/00188). This research was funded by National Science Center Poland, grant number 2018/29/B/NZ2/01010L and 2019/35/B/NZ2/02548.

## Author Contributions

C.W. conceived the project and together with K.A.N and A.S designed the experiments. K.A.N and A.S. performed experiments. A.S. provided significant input on CRISPR/Cas9 experiment and performed microinjections. N.O. and B.K. coordinated single cell encapsulation and library construction. K.A.N. performed sequencing, bioinformatic and statistical analysis. K.A.N designed and developed R shiny app. M.M. contributed to data analysis and R shiny app. R.H., and M.Z. performed spatial analysis. K.A.N. performed morphological analysis. C.W., K.A.N., and M.Z. wrote the manuscript. All authors have read and approved the final paper.

## Notes

### Competing Interest Statement

The authors have declared no competing interest.

## References

1. Harvey RP. Patterning the vertebrate heart. Nat. Rev. Genet. 2002;3:544–56.

2. Samuels MA. The Brain–Heart Connection. Circulation 2007;116:77–84.

3. Moorman AF, de Jong F, Denyn MM, Lamers WH. Development of the cardiac conduction system. Circ. Res. 1998;82:629–44.

4. Sugi Y, Markwald RR. Formation and Early Morphogenesis of Endocardial Endothelial Precursor Cells and the Role of Endoderm. Dev. Biol. 1996;175:66–83.

5. de Lange FJ, Moorman AFM, Anderson RH, Männer J, Soufan AT, de Gier-de Vries C, et al. Lineage and morphogenetic analysis of the cardiac valves. Circ. Res. 2004;95:645–54.

6. Tian X, Pu WT, Zhou B. Cellular origin and developmental program of coronary angiogenesis. Circ. Res. 2015;116:515–30.

7. Serluca FC. Development of the proepicardial organ in the zebrafish. Dev. Biol. 2008;315:18–27.

8. Tallquist MD, Molkentin JD. Redefining the identity of cardiac fibroblasts. Nat. Rev. Cardiol. 2017;14:484–91.

9. Lugnier C, Meyer A, Charloux A, Andrès E, Gény B, Talha S. The Endocrine Function of the Heart: Physiology and Involvements of Natriuretic Peptides and Cyclic Nucleotide Phosphodiesterases in Heart Failure. J. Clin. Med. 2019;8:1746.

10. Lakhal-Littleton S. Mechanisms of cardiac iron homeostasis and their importance to heart function. Free Radic. Biol. Med. 2019;133:234–7.

11. Staudt D, Stainier D. Uncovering the molecular and cellular mechanisms of heart development using the zebrafish. Annu. Rev. Genet. 2012;46:397–418.

12. Yelon D, Horne SA, Stainier DY. Restricted expression of cardiac myosin genes reveals regulated aspects of heart tube assembly in zebrafish. Dev. Biol. 1999;214:23–37.

13. Stainier DY, Lee RK, Fishman MC. Cardiovascular development in the zebrafish. I. Myocardial fate map and heart tube formation. Dev. Camb. Engl. 1993;119:31–40.

14. Bussmann J, Bakkers J, Schulte-Merker S. Early Endocardial Morphogenesis Requires Scl/Tal1. PLOS Genet. 2007;3:e140.

15. de Pater E, Clijsters L, Marques SR, Lin YF, Garavito-Aguilar ZV, Yelon D, et al. Distinct phases of cardiomyocyte differentiation regulate growth of the zebrafish heart. Dev. Camb. Engl. 2009;136:1633–41.

16. Witzel HR, Cheedipudi S, Gao R, Stainier DYR, Dobreva GD. Isl2b regulates anterior second heart field development in zebrafish. Sci. Rep. 2017;7:41043.

17. Cavanaugh AM, Huang J, Chen JN. Two developmentally distinct populations of neural crest cells contribute to the zebrafish heart. Dev. Biol. 2015;404:103–12.

18. Rocha M, Singh N, Ahsan K, Beiriger A, Prince VE. Neural crest development: Insights from the zebrafish. Dev. Dyn. Off. Publ. Am. Assoc. Anat. 2020;249:88–111.

19. Moriyama Y, Ito F, Takeda H, Yano T, Okabe M, Kuraku S, et al. Evolution of the fish heart by sub/neofunctionalization of an elastin gene. Nat. Commun. 2016;7:10397.

20. Milan DJ, Giokas AC, Serluca FC, Peterson RT, MacRae CA. Notch1b and neuregulin are required for specification of central cardiac conduction tissue. Development 2006;133:1125–32.

21. Arnaout R, Ferrer T, Huisken J, Spitzer K, Stainier DYR, Tristani-Firouzi M, et al. Zebrafish model for human long QT syndrome. Proc. Natl. Acad. Sci. U. S. A. 2007;104:11316–21.

22. Vornanen M, Hassinen M. Zebrafish heart as a model for human cardiac electrophysiology. Channels Austin Tex 2016;10:101–10.

23. Koenig AL, Shchukina I, Amrute J, Andhey PS, Zaitsev K, Lai L, et al. Single-cell transcriptomics reveals cell-type-specific diversification in human heart failure. Nat. Cardiovasc. Res. 2022;1:263– 80.

24. Tucker NR, Chaffin M, Fleming SJ, Hall AW, Parsons VA, Bedi KC, et al. Transcriptional and Cellular Diversity of the Human Heart. Circulation 2020;142:466–82.

25. Litviňuková M, Talavera-López C, Maatz H, Reichart D, Worth CL, Lindberg EL, et al. Cells of the adult human heart. Nature 2020;588:466–72.

26. Asp M, Giacomello S, Larsson L, Wu C, Fürth D, Qian X, et al. A Spatiotemporal Organ-Wide Gene Expression and Cell Atlas of the Developing Human Heart. Cell 2019;179:1647–1660.e19.

27. Jiang M, Xiao Y, E W, Ma L, Wang J, Chen H, et al. Characterization of the Zebrafish Cell Landscape at Single-Cell Resolution. Front. Cell Dev. Biol. [Internet] 2021 [cited 2022 Jul 20];9. Available from: https://www.frontiersin.org/articles/10.3389/fcell.2021.743421

28. Weinberger M, Simões FC, Patient R, Sauka-Spengler T, Riley PR. Functional Heterogeneity within the Developing Zebrafish Epicardium. Dev. Cell 2020;52:574–590.e6.

29. Farnsworth DR, Saunders LM, Miller AC. A single-cell transcriptome atlas for zebrafish development. Dev. Biol. 2020;459:100–8.

30. Honkoop H, de Bakker DE, Aharonov A, Kruse F, Shakked A, Nguyen PD, et al. Single-cell analysis uncovers that metabolic reprogramming by ErbB2 signaling is essential for cardiomyocyte proliferation in the regenerating heart. eLife 2019;8:e50163.

31. Burkhard SB, Bakkers J. Spatially resolved RNA-sequencing of the embryonic heart identifies a role for Wnt/β-catenin signaling in autonomic control of heart rate. eLife 2018;7:e31515.

32. Minhas R, Loeffler-Wirth H, Siddiqui YH, Obrębski T, Vashisht S, Nahia KA, et al. Transcriptome profile of the sinoatrial ring reveals conserved and novel genetic programs of the zebrafish pacemaker. BMC Genomics 2021;22:715.

33. Abu Nahia K, Migdał M, Quinn TA, Poon KL, Łapiński M, Sulej A, et al. Genomic and physiological analyses of the zebrafish atrioventricular canal reveal molecular building blocks of the secondary pacemaker region. Cell. Mol. Life Sci. 2021;78:6669–87.

34. Ohtani K, Suzuki Y, Eda S, Kawai T, Kase T, Yamazaki H, et al. Molecular Cloning of a Novel Human Collectin from Liver (CL-L1)*. J. Biol. Chem. 1999;274:13681–9.

35. Malik N, Canfield VA, Beckers MC, Gros P, Levenson R. Identification of the Mammalian Na,K-ATPase β3 Subunit *. J. Biol. Chem. 1996;271:22754–8.

36. Poon KL, Liebling M, Kondrychyn I, Garcia-Lecea M, Korzh V. Zebrafish cardiac enhancer trap lines: new tools for in vivo studies of cardiovascular development and disease. Dev. Dyn. Off. Publ. Am. Assoc. Anat. 2010;239:914–26.

37. Poon KL, Liebling M, Kondrychyn I, Brand T, Korzh V. Development of the cardiac conduction system in zebrafish. Gene Expr. Patterns GEP 2016;21:89–96.

38. Rohr S, Otten C, Abdelilah-Seyfried S. Asymmetric involution of the myocardial field drives heart tube formation in zebrafish. Circ. Res. 2008;102:e12–19.

39. Huang CJ, Tu CT, Hsiao CD, Hsieh FJ, Tsai HJ. Germ-line transmission of a myocardium-specific GFP transgene reveals critical regulatory elements in the cardiac myosin light chain 2 promoter of zebrafish. Dev. Dyn. 2003;228:30–40.

40. D’Amico L, Scott IC, Jungblut B, Stainier DYR. A Mutation in Zebrafish hmgcr1b Reveals a Role for Isoprenoids in Vertebrate Heart-Tube Formation. Curr. Biol. 2007;17:252–9.

41. Kimmel CB, Ballard WW, Kimmel SR, Ullmann B, Schilling TF. Stages of embryonic development of the zebrafish. Dev. Dyn. Off. Publ. Am. Assoc. Anat. 1995;203:253–310.

42. Karlsson J, von Hofsten J, Olsson PE. Generating transparent zebrafish: a refined method to improve detection of gene expression during embryonic development. Mar. Biotechnol. N. Y. N 2001;3:522–7.

43. Lombardo VA, Otten C, Abdelilah-Seyfried S. Large-scale zebrafish embryonic heart dissection for transcriptional analysis. J. Vis. Exp. JoVE 2015;52087.

44. Bresciani E, Broadbridge E, Liu PP. An efficient dissociation protocol for generation of single cell suspension from zebrafish embryos and larvae. MethodsX 2018;5:1287–90.

45. Zheng GXY, Terry JM, Belgrader P, Ryvkin P, Bent ZW, Wilson R, et al. Massively parallel digital transcriptional profiling of single cells. Nat. Commun. 2017;8:14049.

46. Satija R, Farrell JA, Gennert D, Schier AF, Regev A. Spatial reconstruction of single-cell gene expression data. Nat. Biotechnol. 2015;33:495–502.

47. R Core Team. R: a language and environment for statistical computing [Internet]. 2019 [cited 2021 Jan 7];Available from: https://www.gbif.org/tool/81287/r-a-language-and-environment-for-statistical-computing

48. Osorio D, Cai JJ. Systematic determination of the mitochondrial proportion in human and mice tissues for single-cell RNA-sequencing data quality control. Bioinforma. Oxf. Engl. 2021;37:963– 7.

49. McGinnis CS, Murrow LM, Gartner ZJ. DoubletFinder: Doublet Detection in Single-Cell RNA Sequencing Data Using Artificial Nearest Neighbors. Cell Syst. 2019;8:329–337.e4.

50. Hafemeister C, Satija R. Normalization and variance stabilization of single-cell RNA-seq data using regularized negative binomial regression. Genome Biol. 2019;20:296.

51. Stuart T, Butler A, Hoffman P, Hafemeister C, Papalexi E, Mauck WM, et al. Comprehensive Integration of Single-Cell Data. Cell 2019;177:1888–1902.e21.

52. Mi H, Muruganujan A, Ebert D, Huang X, Thomas PD. PANTHER version 14: more genomes, a new PANTHER GO-slim and improvements in enrichment analysis tools. Nucleic Acids Res. 2019;47:D419–26.

53. Yu G, Wang LG, Han Y, He QY. clusterProfiler: an R Package for Comparing Biological Themes Among Gene Clusters. OMICS J. Integr. Biol. 2012;16:284–7.

54. Chloe Li KY, Cook AC, Lovering RC. GOing Forward With the Cardiac Conduction System Using Gene Ontology. Front. Genet. [Internet] 2022 [cited 2023 Apr 27];13. Available from: https://www.frontiersin.org/articles/10.3389/fgene.2022.802393

55. Chodkowski M, Zieleziński A, Anbalagan S. A ligand-receptor interactome atlas of the zebrafish [Internet]. 2023 [cited 2023 Apr 14];2022.12.15.520415. Available from: https://www.biorxiv.org/content/10.1101/2022.12.15.520415v2

56. Labun K, Montague TG, Krause M, Torres Cleuren YN, Tjeldnes H, Valen E. CHOPCHOP v3: expanding the CRISPR web toolbox beyond genome editing. Nucleic Acids Res. 2019;47:W171– 4.

57. Kroll F, Powell GT, Ghosh M, Gestri G, Antinucci P, Hearn TJ, et al. A simple and effective F0 knockout method for rapid screening of behaviour and other complex phenotypes. eLife 2021;10:e59683.

58. Burger A, Lindsay H, Felker A, Hess C, Anders C, Chiavacci E, et al. Maximizing mutagenesis with solubilized CRISPR-Cas9 ribonucleoprotein complexes. Dev. Camb. Engl. 2016;143:2025– 37.

59. Wu RS, Lam II, Clay H, Duong DN, Deo RC, Coughlin SR. A Rapid Method for Directed Gene Knockout for Screening in G0 Zebrafish. Dev. Cell 2018;46:112–125.e4.

60. Stainier DY, Fouquet B, Chen JN, Warren KS, Weinstein BM, Meiler SE, et al. Mutations affecting the formation and function of the cardiovascular system in the zebrafish embryo. Dev. Camb. Engl. 1996;123:285–92.

61. Sehnert AJ, Huq A, Weinstein BM, Walker C, Fishman M, Stainier DYR. Cardiac troponin T is essential in sarcomere assembly and cardiac contractility. Nat. Genet. 2002;31:106–10.

62. Lyons MS, Bell B, Stainier D, Peters KG. Isolation of the zebrafish homologues for the tie-1 and tie-2 endothelium-specific receptor tyrosine kinases. Dev. Dyn. Off. Publ. Am. Assoc. Anat. 1998;212:133–40.

63. Gering M, Rodaway AR, Göttgens B, Patient RK, Green AR. The SCL gene specifies haemangioblast development from early mesoderm. EMBO J. 1998;17:4029–45.

64. Kim CH, Ueshima E, Muraoka O, Tanaka H, Yeo SY, Huh TL, et al. Zebrafish elav/HuC homologue as a very early neuronal marker. Neurosci. Lett. 1996;216:109–12.

65. Das A, Crump JG. Bmps and Id2a Act Upstream of Twist1 To Restrict Ectomesenchyme Potential of the Cranial Neural Crest. PLoS Genet. 2012;8:e1002710.

66. Li M, Zhao C, Wang Y, Zhao Z, Meng A. Zebrafish sox9b is an early neural crest marker. Dev. Genes Evol. 2002;212:203–6.

67. Simões-Costa M, Bronner ME. Insights into neural crest development and evolution from genomic analysis. Genome Res. 2013;23:1069–80.

68. Kollmar R, Nakamura SK, Kappler JA, Hudspeth AJ. Expression and phylogeny of claudins in vertebrate primordia. Proc. Natl. Acad. Sci. 2001;98:10196–201.

69. Dubois GM, Haftek Z, Crozet C, Garrone R, Le Guellec D. Structure and spatio temporal expression of the full length DNA complementary to RNA coding for α2 type I collagen of zebrafish. Gene 2002;294:55–65.

70. Caputo L, Witzel HR, Kolovos P, Cheedipudi S, Looso M, Mylona A, et al. The Isl1/Ldb1 Complex Orchestrates Genome-wide Chromatin Organization to Instruct Differentiation of Multipotent Cardiac Progenitors. Cell Stem Cell 2015;17:287–99.

71. Sumanas S, Zhang B, Dai R, Lin S. 15-zinc finger protein Bloody Fingers is required for zebrafish morphogenetic movements during neurulation. Dev. Biol. 2005;283:85–96.

72. Thompson MA, Ransom DG, Pratt SJ, MacLennan H, Kieran MW, Detrich HW, et al. The cloche and spadetail genes differentially affect hematopoiesis and vasculogenesis. Dev. Biol. 1998;197:248–69.

73. Brownlie A, Donovan A, Pratt SJ, Paw BH, Oates AC, Brugnara C, et al. Positional cloning of the zebrafish sauternes gene: a model for congenital sideroblastic anaemia. Nat. Genet. 1998;20:244– 50.

74. Lieschke GJ, Oates AC, Crowhurst MO, Ward AC, Layton JE. Morphologic and functional characterization of granulocytes and macrophages in embryonic and adult zebrafish. Blood 2001;98:3087–96.

75. Qian F, Zhen F, Ong C, Jin SW, Meng Soo H, Stainier DYR, et al. Microarray analysis of zebrafish cloche mutant using amplified cDNA and identification of potential downstream target genes. Dev. Dyn. Off. Publ. Am. Assoc. Anat. 2005;233:1163–72.

76. Zakrzewska A, Cui C, Stockhammer OW, Benard EL, Spaink HP, Meijer AH. Macrophage-specific gene functions in Spi1-directed innate immunity. Blood 2010;116:e1–11.

77. Lin HF, Traver D, Zhu H, Dooley K, Paw BH, Zon LI, et al. Analysis of thrombocyte development in CD41-GFP transgenic zebrafish. Blood 2005;106:3803–10.

78. Beis D, Bartman T, Jin SW, Scott IC, D’Amico LA, Ober EA, et al. Genetic and cellular analyses of zebrafish atrioventricular cushion and valve development. Dev. Camb. Engl. 2005;132:4193– 204.

79. Fish JE, Wythe JD, Xiao T, Bruneau BG, Stainier DYR, Srivastava D, et al. A Slit/miR-218/Robo regulatory loop is required during heart tube formation in zebrafish. Dev. Camb. Engl. 2011;138:1409–19.

80. Dries R, Lange A, Heiny S, Berghaus KI, Bastmeyer M, Bentrop J. Cell Proliferation and Collective Cell Migration During Zebrafish Lateral Line System Development Are Regulated by Ncam/Fgf-Receptor Interactions. Front. Cell Dev. Biol. [Internet] 2021 [cited 2023 Apr 18];8. Available from: https://www.frontiersin.org/articles/10.3389/fcell.2020.591011

81. Ieda M, Kanazawa H, Kimura K, Hattori F, Ieda Y, Taniguchi M, et al. Sema3a maintains normal heart rhythm through sympathetic innervation patterning. Nat. Med. 2007;13:604–12.

82. Vahouny GV, Tamboli A, Albert EN, Weglicki WB, Spanier AM. Structural and functional characteristics of cardiac myocytes. Basic Res. Cardiol. 1985;80 Suppl 2:45–9.

83. Berdougo E, Coleman H, Lee DH, Stainier DYR, Yelon D. Mutation of weak atrium/atrial myosin heavy chain disrupts atrial function and influences ventricular morphogenesis in zebrafish. Dev. Camb. Engl. 2003;130:6121–9.

84. Yelon D, Stainier DYR. Patterning during organogenesis: genetic analysis of cardiac chamber formation. Semin. Cell Dev. Biol. 1999;10:93–8.

85. Gladka MM, Molenaar B, de Ruiter H, van der Elst S, Tsui H, Versteeg D, et al. Single-Cell Sequencing of the Healthy and Diseased Heart Reveals Cytoskeleton-Associated Protein 4 as a New Modulator of Fibroblasts Activation. Circulation 2018;138:166–80.

86. Liang D, Xue J, Geng L, Zhou L, Lv B, Zeng Q, et al. Cellular and molecular landscape of mammalian sinoatrial node revealed by single-cell RNA sequencing. Nat. Commun. 2021;12:287.

87. Stoyek MR, Quinn TA, Croll RP, Smith FM. Zebrafish heart as a model to study the integrative autonomic control of pacemaker function. Am. J. Physiol.-Heart Circ. Physiol. 2016;311:H676–88.

88. Tessadori F, van Weerd JH, Burkhard SB, Verkerk AO, de Pater E, Boukens BJ, et al. Identification and functional characterization of cardiac pacemaker cells in zebrafish. PloS One 2012;7:e47644.

89. Haack T, Abdelilah-Seyfried S. The force within: endocardial development, mechanotransduction and signalling during cardiac morphogenesis. Dev. Camb. Engl. 2016;143:373–86.

90. Kim JD, Kim HJ, Koun S, Ham HJ, Kim MJ, Rhee M, et al. Zebrafish Crip2 Plays a Critical Role in Atrioventricular Valve Development by Downregulating the Expression of ECM Genes in the Endocardial Cushion. Mol. Cells 2014;37:406–11.

91. Schwarze U, Hata RI, McKusick VA, Shinkai H, Hoyme HE, Pyeritz RE, et al. Rare Autosomal Recessive Cardiac Valvular Form of Ehlers-Danlos Syndrome Results from Mutations in the COL1A2 Gene That Activate the Nonsense-Mediated RNA Decay Pathway. Am. J. Hum. Genet. 2004;74:917–30.

92. Delvaeye M, De Vriese A, Zwerts F, Betz I, Moons M, Autiero M, et al. Role of the 2 zebrafish survivingenes in vasculo-angiogenesis, neurogenesis, cardiogenesis and hematopoiesis. BMC Dev. Biol. 2009;9:25.

93. Gomez GA, Veldman MB, Zhao Y, Burgess S, Lin S. Discovery and Characterization of Novel Vascular and Hematopoietic Genes Downstream of Etsrp in Zebrafish. PLOS ONE 2009;4:e4994.

94. Chen E, Hermanson S, Ekker SC. Syndecan-2 is essential for angiogenic sprouting during zebrafish development. Blood 2004;103:1710–9.

95. Zhao B, Chun C, Liu Z, Horswill MA, Pramanik K, Wilkinson GA, et al. Nogo-B receptor is essential for angiogenesis in zebrafish via Akt pathway. Blood 2010;116:5423–33.

96. Pociute K, Schumacher JA, Sumanas S. Clec14a genetically interacts with Etv2 and Vegf signaling during vasculogenesis and angiogenesis in zebrafish. BMC Dev. Biol. 2019;19:6.

97. Sun G, Liu K, Wang X, Liu X, He Q, Hsiao CD. Identification and Expression Analysis of Zebrafish (Danio rerio) E-Selectin during Embryonic Development. Molecules 2015;20:18539–50.

98. Cao G, O’Brien CD, Zhou Z, Sanders SM, Greenbaum JN, Makrigiannakis A, et al. Involvement of human PECAM-1 in angiogenesis and in vitro endothelial cell migration. Am. J. Physiol. Cell Physiol. 2002;282:C1181–1190.

99. Isogai C, Laug WE, Shimada H, Declerck PJ, Stins MF, Durden DL, et al. Plasminogen Activator Inhibitor-1 Promotes Angiogenesis by Stimulating Endothelial Cell Migration toward Fibronectin1. Cancer Res. 2001;61:5587–94.

100. Tang J, Hu G, Hanai J ichi, Yadlapalli G, Lin Y, Zhang B, et al. A Critical Role for Calponin 2 in Vascular Development *. J. Biol. Chem. 2006;281:6664–72.

101. Fedele L, Brand T. The Intrinsic Cardiac Nervous System and Its Role in Cardiac Pacemaking and Conduction. J. Cardiovasc. Dev. Dis. 2020;7:54.

102. Sato A, Takeda H. Neuronal Subtypes Are Specified by the Level of neurod Expression in the Zebrafish Lateral Line. J. Neurosci. 2013;33:556–62.

103. Veldman MB, Bemben MA, Goldman D. Tuba1a gene expression is regulated by KLF6/7 and is necessary for CNS development and regeneration in zebrafish. Mol. Cell. Neurosci. 2010;43:370– 83.

104. Oehlmann VD, Berger S, Sterner C, Korsching SI. Zebrafish beta tubulin 1 expression is limited to the nervous system throughout development, and in the adult brain is restricted to a subset of proliferative regions. Gene Expr. Patterns 2004;4:191–8.

105. Milanese C, Sager JJ, Bai Q, Farrell TC, Cannon JR, Greenamyre JT, et al. Hypokinesia and Reduced Dopamine Levels in Zebrafish Lacking β- and γ1-Synucleins*. J. Biol. Chem. 2012;287:2971–83.

106. Winkler C, Schafer M, Duschl J, Schartl M, Volff JN. Functional divergence of two zebrafish midkine growth factors following fish-specific gene duplication. Genome Res. 2003;13:1067–81.

107. Wu MY, Ramel MC, Howell M, Hill CS. SNW1 is a critical regulator of spatial BMP activity, neural plate border formation, and neural crest specification in vertebrate embryos. PLoS Biol. 2011;9:e1000593.

108. Knight RD, Nair S, Nelson SS, Afshar A, Javidan Y, Geisler R, et al. lockjaw encodes a zebrafish tfap2a required for early neural crest development. Dev. Camb. Engl. 2003;130:5755–68.

109. Pöpperl H, Rikhof H, Chang H, Haffter P, Kimmel CB, Moens CB. lazarus is a novel pbx gene that globally mediates hox gene function in zebrafish. Mol. Cell 2000;6:255–67.

110. Tanaka H, Yoshida T, Miyamoto N, Motoike T, Kurosu H, Shibata K, et al. Characterization of a family of endogenous neuropeptide ligands for the G protein-coupled receptors GPR7 and GPR8. Proc. Natl. Acad. Sci. U. S. A. 2003;100:6251–6.

111. Van Camp KA, Baggerman G, Blust R, Husson SJ. Peptidomics of the zebrafish Danio rerio: In search for neuropeptides. J. Proteomics 2017;150:290–6.

112. Söderberg C, Wraith A, Ringvall M, Yan YL, Postlethwait JH, Brodin L, et al. Zebrafish Genes for Neuropeptide Y and Peptide YY Reveal Origin by Chromosome Duplication from an Ancestral Gene Linked to the Homeobox Cluster. J. Neurochem. 2000;75:908–18.

113. Devos N, Deflorian G, Biemar F, Bortolussi M, Martial JA, Peers B, et al. Differential expression of two somatostatin genes during zebrafish embryonic development. Mech. Dev. 2002;115:133–7.

114. Roth LWA, Bormann P, Bonnet A, Reinhard E. β-Thymosin is required for axonal tract formation in developing zebrafish brain. Development 1999;126:1365–74.

115. Ma M, Ramirez AD, Wang T, Roberts RL, Harmon KE, Schoppik D, et al. Zebrafish dscaml1 Deficiency Impairs Retinal Patterning and Oculomotor Function. J. Neurosci. Off. J. Soc. Neurosci. 2020;40:143–58.

116. Gao X, Metzger U, Panza P, Mahalwar P, Alsheimer S, Geiger H, et al. A Floor-Plate Extracellular Protein-Protein Interaction Screen Identifies Draxin as a Secreted Netrin-1 Antagonist. Cell Rep. 2015;12:694–708.

117. Tongiorgi E, Bernhardt RR, Zinn K, Schachner M. Tenascin-C mRNA is expressed in cranial neural crest cells, in some placodal derivatives, and in discrete domains of the embryonic zebrafish brain. J. Neurobiol. 1995;28:391–407.

118. Sawai S, Campos-Ortega JA. A zebrafish Id homologue and its pattern of expression during embryogenesis. Mech. Dev. 1997;65:175–85.

119. Cera AA, Cacci E, Toselli C, Cardarelli S, Bernardi A, Gioia R, et al. Egr-1 Maintains NSC Proliferation and Its Overexpression Counteracts Cell Cycle Exit Triggered by the Withdrawal of Epidermal Growth Factor. Dev. Neurosci. 2018;40:223–33.

120. Hortells L, Meyer EC, Thomas ZM, Yutzey KE. Periostin-expressing Schwann cells and endoneurial cardiac fibroblasts contribute to sympathetic nerve fasciculation after birth. J. Mol. Cell. Cardiol. 2021;154:124–36.

121. Chi NC, Shaw RM, Jungblut B, Huisken J, Ferrer T, Arnaout R, et al. Genetic and Physiologic Dissection of the Vertebrate Cardiac Conduction System. PLOS Biol. 2008;6:e109.

122. Sylvén C, Wärdell E, Månsson-Broberg A, Cingolani E, Ampatzis K, Larsson L, et al. High cardiomyocyte diversity in human early prenatal heart development. iScience 2023;26:105857.

123. Frey N, Richardson JA, Olson EN. Calsarcins, a novel family of sarcomeric calcineurin-binding proteins. Proc. Natl. Acad. Sci. 2000;97:14632–7.

124. Harrison MRM, Bussmann J, Huang Y, Zhao L, Osorio A, Burns CG, et al. Chemokine guided angiogenesis directs coronary vasculature formation in zebrafish. Dev. Cell 2015;33:442–54.

125. Ks W, K R, S PD, V K, W H, Jd U, et al. Hedgehog signaling is required for differentiation of endocardial progenitors in zebrafish. Dev. Biol. [Internet] 2012 [cited 2023 Apr 18];361. Available from: https://pubmed.ncbi.nlm.nih.gov/22119054/

126. Sánchez-Iranzo H, Galardi-Castilla M, Sanz-Morejón A, González-Rosa JM, Costa R, Ernst A, et al. Transient fibrosis resolves via fibroblast inactivation in the regenerating zebrafish heart. Proc. Natl. Acad. Sci. U. S. A. 2018;115:4188–93.

127. Hu B, Lelek S, Spanjaard B, El-Sammak H, Simões MG, Mintcheva J, et al. Origin and function of activated fibroblast states during zebrafish heart regeneration. Nat. Genet. 2022;54:1227–37.

128. McDonough AA, Zhang Y, Shin V, Frank JS. Subcellular distribution of sodium pump isoform subunits in mammalian cardiac myocytes. Am. J. Physiol.-Cell Physiol. 1996;270:C1221–7.

129. Tulloch LB, Howie J, Wypijewski KJ, Wilson CR, Bernard WG, Shattock MJ, et al. The inhibitory effect of phospholemman on the sodium pump requires its palmitoylation. J. Biol. Chem. 2011;286:36020–31.

130. Munye MM, Diaz-Font A, Ocaka L, Henriksen ML, Lees M, Brady A, et al. COLEC10 is mutated in 3MC patients and regulates early craniofacial development. PLoS Genet. 2017;13:e1006679.

