## Supplementary material for "scRNA-seq reveals the diversity of the developing cardiac cell lineage and molecular building blocks of the primary pacemaker": Supp.Figures

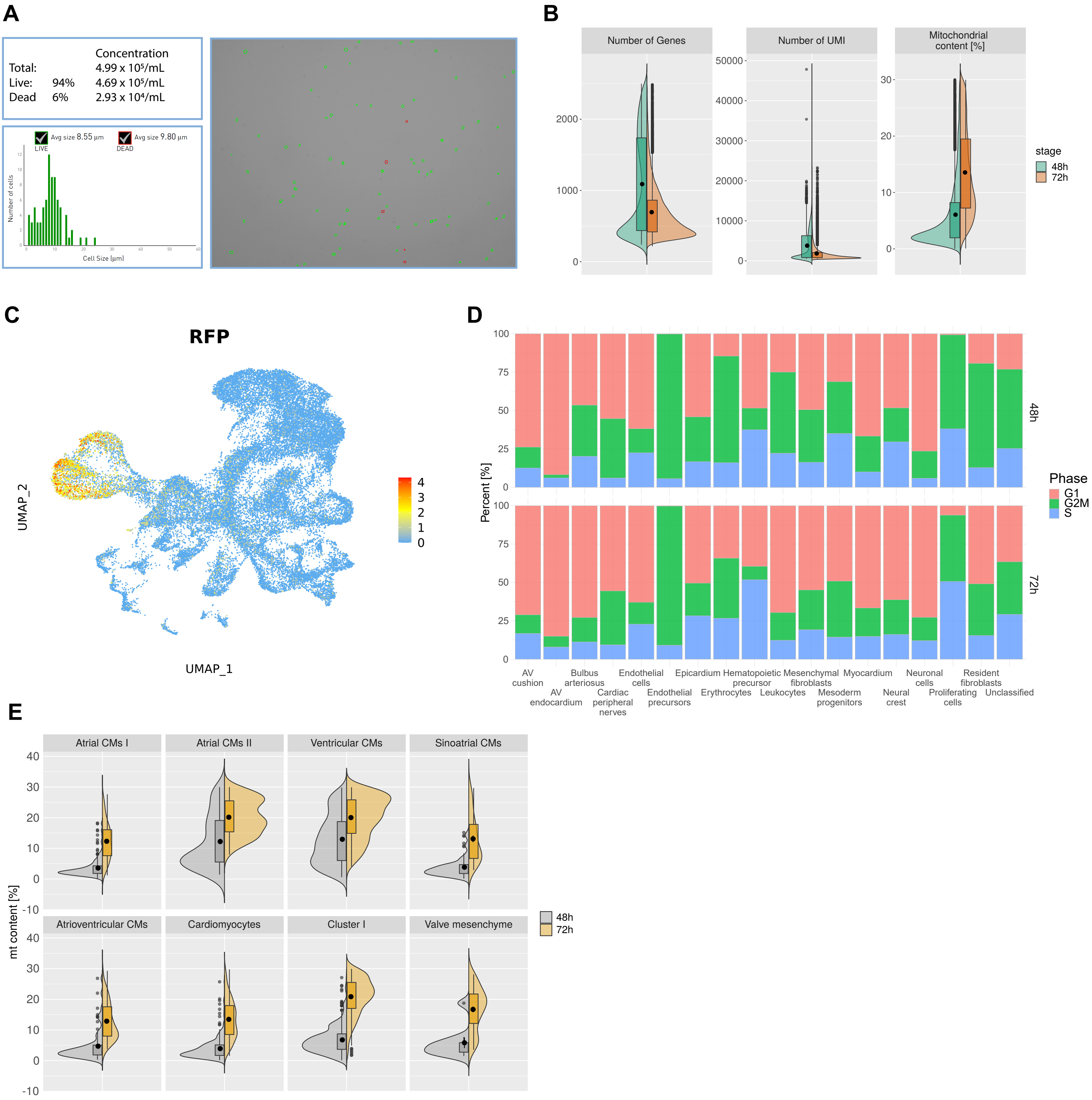


**Supplementary Figure 1. Quality parameters of scRNA-seq dataset. A** Quality and quantity check of single cell suspension by automated cell counter. Viable cells are marked as green. **B** Violin plot showing mitochondrial gene content, number of expressed genes, and UMI at each developmental stage. **C** UMAP plot of cardiac cell clusters indicating the cardiomyocyte expression of RFP reporter gene. **D** Composition of cells in each main clusters expressed as a percentage according to their cell cycle phase. **E** Mitochondrial gene content in each myocardial cluster depending on developmental stage.

**A**

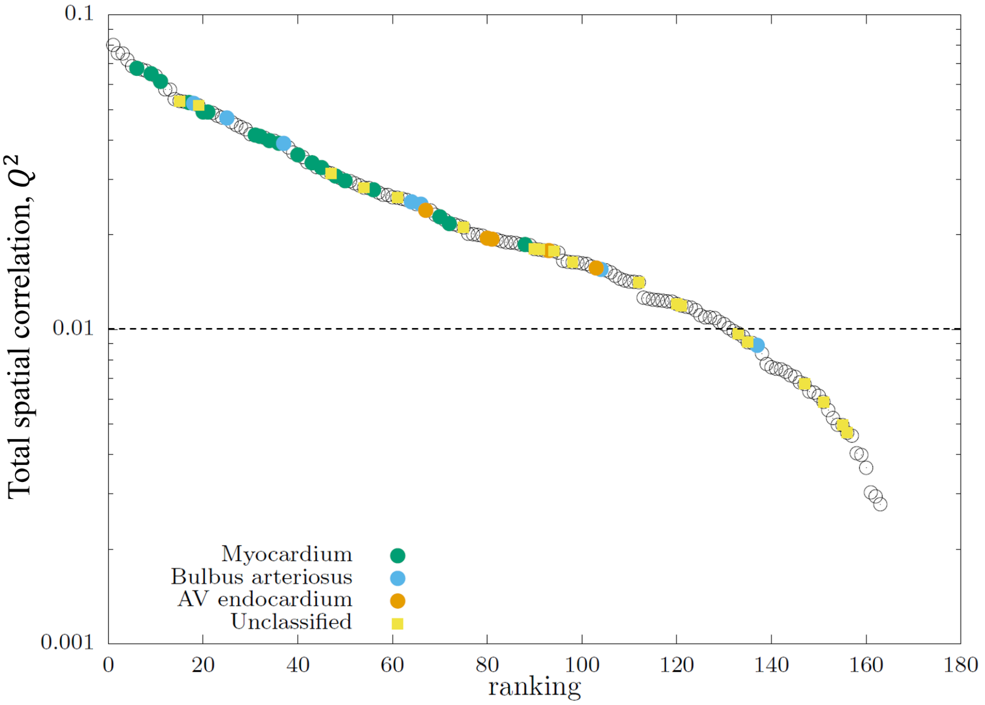


| Cluster | $Q^2$ |
|---|---|
| Bulbus arteriosus | 0,1070 |
| Erythrocytes | 0,0830 |
| Mesoderm progenitors | 0,0760 |
| Resident fibroblasts | 0,0610 |
| Proliferating cells | 0,0520 |
| Epicardium | 0,0450 |
| Hematopoietic precursor | 0,0350 |
| Endothelial precursors | 0,0300 |
| Myocardium | 0,0300 |
| AV endocardium | 0,0210 |
| Endothelial cells | 0,0200 |
| Leukocytes | 0,0190 |
| AV cushion | 0,0180 |
| Neuronal cells | 0,0160 |
| Cardiac peripheral nerves | 0,0150 |
| Neural crest | 0,0130 |
| Mesenchymal fibroblasts | 0,0130 |
| Unclassified | 0,0050 |

**B**

| Cluster | Composition |
|---|---|
| 115 | 99.29% Mesoderm progenitors, 0.71% Bulbus arteriosus |
| 26 | 100% Mesoderm progenitors |
| 31 | 95.21% Erythrocytes, 4.79% Unclassified |
| 28 | 100% Erythrocytes |
| 5 | 86.67% Bulbus arteriosus, 2.86% Epicardium, 10.48% Unclassified |

**Supplementary Figure 2. Correlations between main and fine-grained clusters. A** Spatial correlation between main cluster and 164 fine-grained clusters. $Q^2$ represents the sum of the square of the Pearson correlation coefficient from all sections normalised by the number of sections. The $Q^2$ for main clusters is given in the table. The $Q^2$ for fine-grained clusters was ranked from the highest to the lowest. The highlighted fine-grained clusters are associated with one of the selected main cluster (“Myocardium”, “Bulbus arteriosus”, “AV endocardium”, and “Unclassified”), if 90% of points in that cluster belong to the respective main cluster. The “Myocardium” fine-grained clusters are quite high with “Bulbus arteriosus” fine-grained clusters ranked slightly lower. The dashed line indicates roughly the maximum level of $Q^2$ obtained by calculating spatial correlation with randomized gene order for several randomly selected fine-grained clusters. **B** Top 5 fine-grained clusters with the highest $Q^2$.

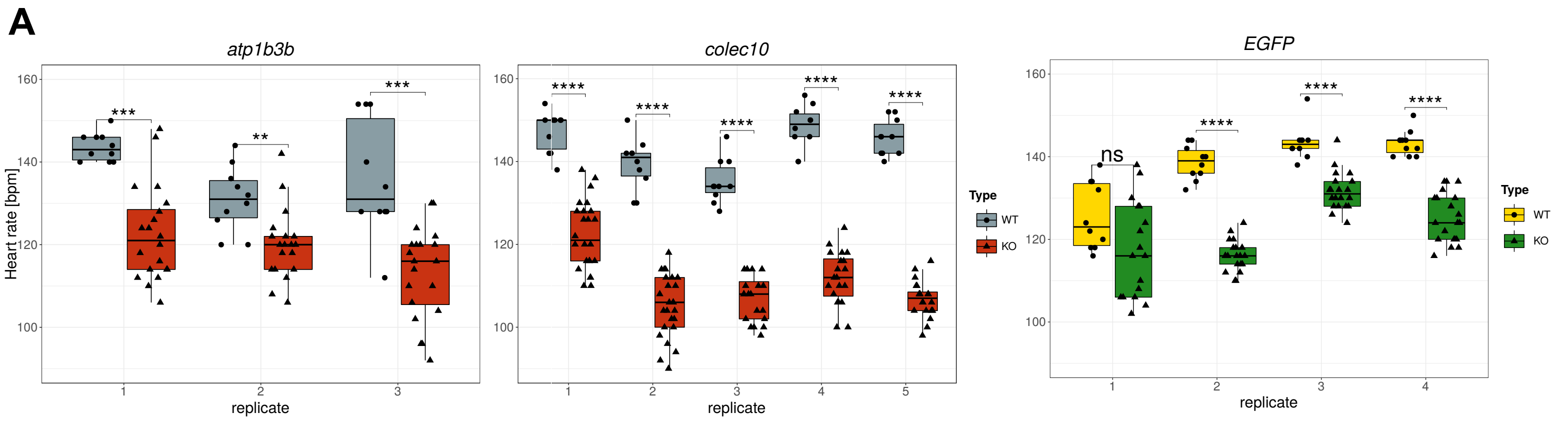


**Supplementary Figure 3. Differences in heartbeat rate in each replicate of targeted genes. A** Boxplot showing differences in heart rate between each targeted gene knockout embryos versus corresponding uninjected control siblings (individual replicates). Statistical test: Wilcoxon rank sum test.
