## Supplementary material for "scRNA-seq reveals the diversity of the developing cardiac cell lineage and molecular building blocks of the primary pacemaker": Supp.Methods

**Gene expression and spatial cluster correlation analysis**

The gene expression correlation $\rho$genes for main clusters using fine-grained clusters was calculated by applying the following steps: (1) ENSEMBL gene IDs were used to match the data from the main and fine-grained clusters, (2) in the cluster data, the log2FC were converted to fold change values, (3) the prediction of gene values for the main cluster is made as the weighted average of gene values (weighted by the percentage of the fine grained cluster that is in each main cluster, see Supplementary Table 6) of this particular cluster fine-grained constituents. (4) the Pearson correlation coefficient $\rho$genes between the predicted gene values and actual gene values were estimated. The gene expression correlations $\rho$genes for fine-grained clusters using main clusters is calculated analogously to the above procedure, but in step (3) the prediction of gene values for the fine-grained cluster is given by the weighted average of gene values of main clusters that constitute this particular fine-grained cluster. The spatial cluster correlation $\rho$spatial for main clusters using fine-grained clusters was calculated by applying the following steps: (1) the spatial correlations $\rho(x)$ were calculated for main and fine-grained clusters (see Spatial correlations); where $x$ is section number, (2) the prediction of $\rho(x)$ for the main cluster is given as the weighted average of $\rho(x)$ of this particular cluster fine-grained constituents. (3) the Pearson correlation coefficient between the predicted values and actual values is estimated whilst ignoring the last three sections. Similarly, the spatial cluster correlations $\rho$spatial for fine-grained clusters using main clusters are calculated analogously to the above procedure, but in step (2) the decomposition of a fine-grained cluster into main cluster constituents is used.

The cluster “Myocardium” was negatively correlated with sections representing the venous pole which are known to be largely devoid of this cell type [19], while positively correlated with subsequent sections representing the cardiac chambers. In contrast, the cluster “Bulbus arteriosus” exhibited a complementary correlation pattern, which reflected their localization at the venous pole. For clusters representing cells of the atrioventricular canal, “AV endocardium” and “AV cushion”, the correlation is highest with sections 20-30 which agrees with the intermediary position of this structure along the heart tube. The clusters “Neural crest”, “cardiac peripheral nerves”, and “neuronal cells” showed a high correlation with sections representing the outflow, atrioventricular canal, and inflow region, in line with previous reports in mammals [78,79].

We took advantage of then asked whether we could utilize spatial correlation to interrogate the components of each main cluster at a higher resolution. We reasoned that if the main cluster is composed of homogenous cell types, they should possess a similar level of correlation between them, as well as with the main cluster as a whole. To this end, we applied the same spatial correlation analysis to 164 fine-grained clusters (resolution = 15) and the respective main clusters as reference. The fine-grained clusters associated with “Myocardium” showed the highest correlation, while those of the “Bulbus arteriosus” ranked slightly lower (Supplementary Fig. 2A). The top 5 fine-grained clusters with the highest spatial correlation include cells originating from “mesoderm progenitors” (Supplementary Fig. 2B), which had the highest correlation with the outflow region. The main clusters overall exhibit high levels of both gene and spatial cluster correlation, except for the “unclassified” cluster which was heterogenous in both spatial and gene expression values, and the “mesenchymal fibroblasts” cluster which was homogeneous in terms of gene expression values but spatially heterogeneous (Supplementary Table 6). On the other hand, the fine-grained clusters exhibited high gene expression and spatial correlation (Supplementary Table 6). However, the variability was much higher than for the main clusters. High spatial correlation with low or negative gene correlation for the fine-grained clusters suggested that these clusters might be controlling subparts of the main clusters which have a differentiated role but are otherwise attached to the developmental structure associated with the respective main cluster.
